# Olig2 and Hes regulatory dynamics during motor neuron differentiation revealed by single cell transcriptomics

**DOI:** 10.1101/104307

**Authors:** Andreas Sagner, Zachary B. Gaber, Julien Delile, Jennifer H. Kong, David L. Rousso, Caroline A. Pearson, Steven E. Weicksel, Manuela Melchionda, Neda S. Mousavy Gharavy, James Briscoe, Bennett G. Novitch

## Abstract

During tissue development, multipotent progenitors differentiate into specific cell types in characteristic spatial and temporal patterns. We address the mechanism linking progenitor identity and differentiation rate in the neural tube, where motor neuron (MN) progenitors differentiate more rapidly than other progenitors. Using single cell transcriptomics, we define the transcriptional changes associated with the transition of neural progenitors into MNs. Reconstruction of gene expression dynamics from these data indicate a pivotal role for the MN determinant Olig2 just prior to MN differentiation. Olig2 represses expression of the Notch signaling pathway effectors Hes1 and Hes5. Olig2 repression of Hes5 appears to be direct, via a conserved regulatory element within the Hes5 locus that restricts expression from MN progenitors. These findings reveal a tight coupling between the regulatory networks that control patterning and neuronal differentiation, and demonstrate how Olig2 acts as the developmental pacemaker coordinating the spatial and temporal pattern of MN generation.

## INTRODUCTION

The orderly development of embryonic tissues relies on gene regulatory networks that control patterns of gene expression, tissue growth and cell differentiation (Davidson 2010; Stathopoulos and Levine 2005). Genetic and molecular studies have identified many of the constituents of these networks and have begun to define the regulatory hierarchy between them. Nevertheless, how cell fate assignment is coordinated with proliferation and differentiation remains poorly understood.

An experimentally well-characterised tissue that exemplifies this problem is the vertebrate spinal cord. In ventral regions of the developing spinal cord proliferating progenitors are exposed to a gradient of Sonic Hedgehog (Shh) signalling that controls the expression of a set of homeodomain and basic helix-loop-helix (bHLH) transcription factors (TFs) (Briscoe et al. 2000; Ribes and Briscoe 2009; Briscoe and Ericson 1999). These TFs form a gene regulatory network that progressively allocates progenitor identity, dividing the spinal cord into molecularly discrete domains arrayed along the dorsal-ventral axis (Balaskas et al. 2012; Cohen et al. 2013).This combinatorial transcriptional code determines the subtype identity of the post-mitotic neurons generated by progenitors in each domain, thereby controlling the position at which MNs and interneurons emerge (Briscoe et al. 2000; Jessell 2000; Lee and Pfaff 2001; Alaynick et al. 2011).

Among the first neurons to differentiate in the ventral spinal cord are MNs. In mouse and chick, these are formed over a 2 to 3-day period (Kicheva et al. 2014). During this time, most if not all MN progenitors exit the cell cycle and differentiate, whereas the adjacent progenitor domains that give rise to interneurons continue to divide and consequently differentiate at a much slower pace (Kicheva et al. 2014; Ericson et al. 1992). These differences in differentiation rate play an important role in the elaboration of spinal cord pattern and ensure appropriate numbers of MNs are generated. This raises the question of how the regulatory mechanisms defining MN progenitors prime these cells to differentiate rapidly.

The induction and differentiation of MNs are characterized by a series of gene expression changes. Initially, Shh signaling induces the bHLH protein Olig2, resulting in the repression of the homeodomain protein Irx3 and bHLH protein Bhlhb5, normally expressed in neural progenitors (NPs) dorsal to MNs (Novitch et al. 2001; Zhou and Anderson 2002; Lu et al. 2002; Skaggs et al. 2011). Ectopic expression of Olig2 represses both Irx3 and Bhlhb5, resulting in ectopic MN production (Novitch et al. 2001; Mizuguchi et al. 2001; Skaggs et al. 2011). Conversely, in the absence of *Olig2*, MN generation fails and instead Irx3 and Bhlhb5 expression are maintained and NPs differentiate into ventral interneurons (Zhou and Anderson 2002; Lu et al. 2002; Takebayashi et al. 2002; Skaggs et al. 2011).

The gene regulatory mechanisms that are responsible for the higher rate of neurogenesis of MN progenitors compared to other NPs in the spinal cord are not well understood. Whether Olig2 functions as an activator or inhibitor of neurogenesis is unclear. Initial studies indicated that expression of Olig2 accelerates cell cycle exit (Novitch et al. 2001) and the absence of Olig2 results in a characteristically slower tempo of neuronal differentiation (Zhou and Anderson 2002). Olig2 promotes the expression of the proneural bHLH TF Ngn2 and ectopic expression of Ngn2 causes progenitor cells to exit the cell cycle to differentiate prematurely into neurons (Novitch et al. 2001; Scardigli et al. 2001; Mizuguchi et al. 2001; Bertrand et al. 2002; Lu et al. 2000; Sugimori et al. 2007; Lacomme et al. 2012). These studies also showed that Olig2 acts as a transcriptional repressor to promote Ngn2 expression (Novitch et al. 2001; Mizuguchi et al. 2001), implying that Olig2 promotes Ngn2 expression by negatively regulating the expression of Ngn2 repressors. Candidate Ngn2 repressors include members of the Hairy/Enhancer of Split (Hes) family of transcription factors, which act downstream of the Notch signaling pathway to prevent neuronal differentiation and maintain progenitors in a dividing, undifferentiated state (Ohtsuka et al. 1999; Shimojo et al. 2011; Kageyama et al. 2007).

Although these studies suggested that Olig2 promotes motor neurogenesis, subsequent studies ascribed anti-neurogenic and pro-proliferative functions to Olig2 (reviewed in Meijer et al. 2012). These conclusions were based on the Olig2-mediated repression of the MN marker Hb9 (Mnx1) (Lee et al. 2005) and the cell cycle inhibitor p21 (Ligon et al. 2007), as well as the ability of Olig2 to form heterodimers with Ngn2, which inhibit neurogenic activity (Lee et al. 2005), and the capacity of Olig2 to oppose p53 function (Mehta et al. 2011). Furthermore, addition of Olig2 to TF reprogramming cocktails inhibits reprogramming of fibroblasts to MNs (Son et al. 2011), supporting the idea that Olig2 interferes with the differentiation of MNs. Thus, although the genetic evidence establishes Olig2 as a key determinant of MN identity, the apparently contradictory findings leave unexplained how Olig2 coordinates specification of neuronal identity while determining the rate of differentiation.

Single cell RNA sequencing (scRNA-seq) is emerging as a novel and powerful technology to identify distinct cell types in complex mixtures and to define developmental trajectories during differentiation (Scialdone et al. 2016; Treutlein et al. 2016; Setty et al. 2016; Trapnell et al. 2014; Shin et al. 2015). Here we take advantage of an *in vitro* model that allows the generation of ventral spinal cord cell types from embryonic stem cells (ESCs) to perform scRNA-seq analysis of developing NPs (Gouti et al. 2014). We use these data to reconstruct and validate the differentiation trajectory of MN progenitors, and to infer the gene regulatory mechanisms by which Olig2 promotes MN differentiation. Both *in vivo* and *in vitro* cells commit to MN differentiation asynchronously. This limits the temporal resolution of conventional gene expression assays, potentially obscuring details of the sequence of events during MN differentiation. Here we develop a method to reconstruct the differentiation trajectory from scSEQ data that provides much greater temporal resolution of the transcriptional dynamics during MN differentiation than previously available. This approach identified a sequence of distinct phases in MN differentiation, including two distinct Olig2 expression states. An initial Olig2^LOW^ state, during which Hes1 expression decreases and Olig2 is coexpressed with Hes5, and a subsequent Olig2^HIGH^ state in which high levels of Olig2 promote differentiation by repressing Hes5, thereby indirectly inducing Ngn2. We validate this two-phase model using quantitative image analysis of a fluorescent Olig2 reporter and provide *in vitro* and *in vivo* evidence that Olig2 acts directly on *Hes* genes to promote cell cycle exit and neurogenesis in the MN progenitor domain (pMN domain). Together the data provide a comprehensive view of the regulatory network that controls the specification of MN progenitors and identify a molecular mechanism coordinating the specification of positional identity with differentiation.

## RESULTS

### *In vitro* generation of Motor Neuron and V3 progenitors

To define the sequence of events that lead to the generation of somatic MNs, we took advantage of ESCs, which can be directed to differentiate into spinal NPs *in vitro* (Gouti et al. 2014). This method relies on the exposure of ESCs, cultured as a monolayer, to a brief pulse of Wnt signalling prior to neural induction (Fig 1A). This induces the caudalizing TFs Cdx1,2,4 (Gouti et al. 2014). Subsequently, removal of Wnt signalling and concomitant exposure to retinoic acid (RA) and the Shh signalling agonist SAG results in the generation of NPs expressing progenitor markers characteristic for the ventral spinal cord such as Olig2 and Nkx2.2 (Fig 1B) and MNs expressing postmitotic markers including Islet1 (Isl1), Mnx1 and neuronal class III beta-tubulin (Tubb3) (Fig 1B,C and Fig S1A). These NPs express initially *Hoxb1* and later *Hoxb9* (Fig S1B) and differentiate into *Hoxc6*-positive MNs, characteristic of forelimb level spinal cord MNs (Fig S1B,C) (Dasen et al. 2003; Philippidou and Dasen 2013; Stifani 2014; Gouti et al. 2014).

**Fig 1.**
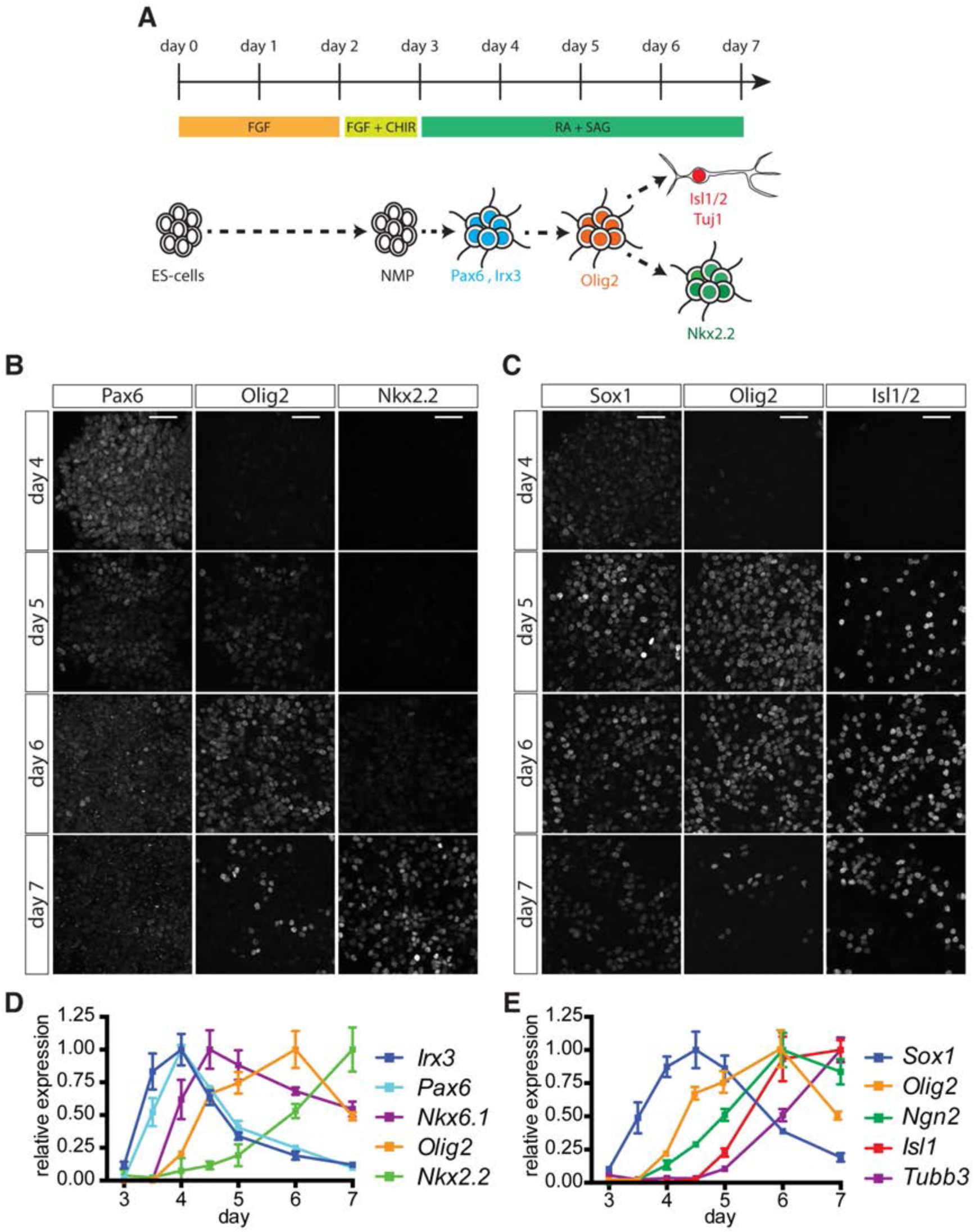
Characterization of MN differentiation from ESCs. (A) Scheme outlining the differentiation protocol. ESCs are plated in N2B27 + FGF for two days, before being exposed to N2B27 + FGF/CHIR, resulting in the production of neuromesodermal progenitors (NMPs) at day 3. Cells are subsequentlyexposedtoretinoicacid(RA)andSAGtopromote differentiation into ventral NPs and MNs. (B, C) Expression of NP (Pax6, Olig2, Nkx2.2, Sox1) and MN (Isl1/2) markers between day 4 to day 7 in differentiating ESCs. (D) RT-qPCR analysis of *Irx3*, *Pax6*, *Nkx6.1*, *Olig2* and *Nkx2.2* expression from day 3 to day 7 reveals progressive ventralization in response to increasing duration of Shh signaling. (E) MN induction after day 5 revealed by RT-qPCR analysis of *Sox1*, *Ngn2*, *Isl1* and *Tubb3*. Scale bars = 40 μm.

*In vivo* NPs respond to both the levels and duration of Shh signalling by transitioning through a succession of progressively more ventral gene expression states (Fig S2A) (Chamberlain et al. 2008; Dessaud et al. 2010; Balaskas et al. 2012; Jeong and McMahon 2005). To further characterize the behaviour of ESCs derived NPs *in vitro,* we first asked whether treatment of NPs with increasing concentrations of SAG (0, 10, 50, 100, 500, and 1000 nM SAG) leads to progressively more ventral cell fates. Generation of NPs in the absence of SAG resulted in the expression of *Pax3*, *Pax7* and *Dbx1*, indicative of a dorsal and intermediate NP identity (Fig S2B). Treatment with 10 nM SAG resulted in the down-regulation of these genes and induction of the pan-ventral marker *Nkx6.1* (Fig S2B,C). Between 50 – 500 nM SAG, expression of the MN progenitor marker *Olig2* was observed, while treatment with 500 and 1000 nM SAG resulted in further ventralization and induction of the p3 determinant *Nkx2.2* (Fig S2B,D). Induction of ventral markers coincided with the successive down-regulation of *Irx3* and *Pax6,* consistent with their *in vivo* expression patterns (Fig S2B,D). Thus, *in vitro* NPs respond to different levels of Shh pathway activity by induction of the same progenitor markers that demarcate NP domains in the embryonic ventral spinal cord.

We next tested whether *in vitro* NPs also displayed progressive ventralization in response to increasing exposure durations to a constant concentration of SAG. To this end, we treated cells with 500 nM SAG from day 3 and quantified gene expression by RT-qPCR over the course of the next few days. At day 3.5, 12 hours after the cessation of Wnt signalling and addition of RA and SAG, cells expressed *Sox1*, *Pax6* and *Irx3*, consistent with the acquisition of NP identity (Fig 1B,D). The absence of ventral markers at this stage indicates that these NPs initially adopt a dorsal/intermediate positional identity (Jeong and McMahon 2005; Dessaud et al. 2010). By day 4, the expression of *Pax6* and *Irx3* were maintained and *Nkx6.1*, which is expressed broadly in the ventral third of the neural tube, was induced (Fig 1B,D). Within 12h of this time point, *Olig2* expression commenced and both *Pax6* and *Irx3* declined (Fig 1B-D). Over the next 48h, *Pax6* and *Irx3* were further repressed, *Nkx2.2* increased and *Olig2* expression began to decline (Fig 1B-D). The order in which these genes were activated and repressed closely resembles the temporal-spatial sequence of progenitor domains in the embryonic spinal cord (Dessaud et al. 2010; Balaskas et al. 2012; Jeong and McMahon 2005), and suggests that under these conditions MN progenitors are generated *in vitro* between day 4.5 and ~day 6.

Consistent with the generation of MN progenitors *in vitro*, *Ngn2* was induced following *Olig2* (Fig 1E); and with a ~12 hours delay we observed markers characteristic of post-mitotic MNs, including *Isl1* and *Tubb3* (Fig 1C,E). Concomitantly, the expression of the NP marker *Sox1* declined (Fig 1C,E). Taken together, these data indicate that this method of directing ESC differentiation recapitulates *in vivo* dynamics of neural tube patterning between approximately e8.5 and e10.5 and results in the production of MN progenitors and MNs characteristic of those normally found at forelimb levels.

### Single cell transcriptome analysis of in vitro NPs

We reasoned that analysing the transcriptome of individual cells would provide insight into the transitions in gene expression associated with the differentiation of MNs and allow the construction of a detailed developmental timeline. We therefore performed scRNA-seq analysis using the Fluidigm-C1 platform on 236 cells isolated from day 4 to day 6 of the differentiation protocol. After applying quality filters (see Analytical Supplement) transcriptomes of 202 cells were retained for subsequent analysis (25 cells from day 4, 68 cells from day 5 and 109 cells from day 6). To identify the cell states present in the dataset, we established a data driven analysis pipeline based on hierarchical clustering and association of gene modules with specific GO terms (see Analytical Supplement). In brief, the data were first filtered by removing genes that did not exceed a Spearman correlation of r>0.4 with at least two other genes (retaining 2287 genes). A combination of hierarchical clustering and automated selection criteria identified 22 gene modules that represent distinct patterns of gene expression across the dataset (see Analytical Supplement and Supplementary File 1). Further functional characterization of these gene modules based on GO terms resulted in the identification of 10 gene modules that were sufficient to assign a cell type classification to each cell in the dataset using hierarchical clustering (Fig S3A and Supplementary File 2). Cells in these clusters showed comparable read counts and number of expressed genes per cell, suggesting that these properties did not bias the clustering (Fig S3B).

Consistent with our previous finding that the spinal NPs generated by differentiation of ESCs share a developmental lineage with trunk mesoderm (Gouti et al. 2014), we observed two mesodermal cell populations in our dataset: paraxial presomitic mesoderm characterized by the expression of *Meox1* and *Foxc1*, and a vascular endothelial population expressing *Dll4* and *Cdh5* (Fig S3A). The remaining cell clusters corresponded to different stages of NPs and differentiating MNs (Fig 2A). Five gene modules were associated with these cells (comprising 306 genes; see Supplementary File 1). Module 1 was enriched for genes upregulated in early NPs, including the TF *Irx3*. Module 2 contained genes expressed in MN progenitors, including the ventral progenitor markers *Olig2* and *Nkx6.1,* and the neural-specific POU TF *Pou3f2* (aka Brn-2). Module 3 comprised a set of genes transiently expressed in MNs as they differentiate, such as the bHLH TFs *Ngn2*, *Neurod1*, *Neurod4* and *Hes6*, the homeodomain TFs *Isl1* and *Lhx3*, and the Notch ligand *Dll1*. Modules 4 and 5 revealed two successive waves of neuronal gene induction. Module 4 contained genes induced early in differentiated MNs such as *Tubb3*, the RNA-binding protein *Elavl3* (aka *HuC*) and the SoxC TF *Sox4*, while Module 5 consisted of genes characteristic of more mature MNs, represented by *Chat* (*Choline acetyltransferase*) and the TFs *Isl2* and *Onecut1* (Velasco et al. 2016; Rhee et al. 2016; Thaler et al. 2004; Tanabe et al. 1998).

**Fig 2.**
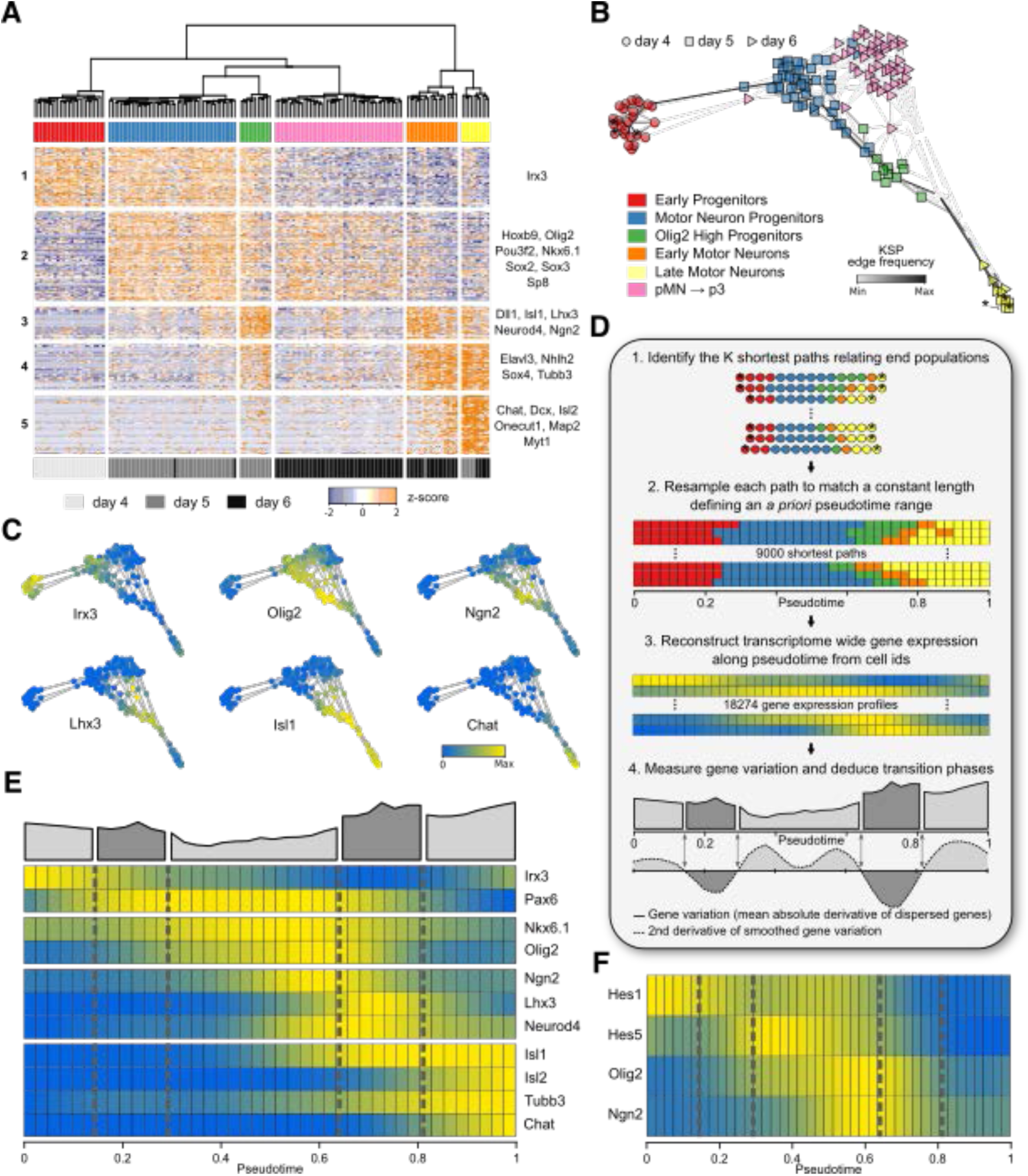
Reconstruction of transcriptional changes during MN differentiation. (A) Identification of NP cell states using hierarchical clustering of gene expression profiles of the individual cells (B) Cell state graph constructed from minimum spanning trees, color coded for the cell populations identified in Fig 2A. Stars indicate start and end cells for the reconstruction of transcriptional changes along pseudotime. Shading of edges between cells indicates how often the edge was used in the reconstruction of gene expression along pseudotime (see Analytical Supplement). (C) Cell state graph color coded for expression levels of *Irx3*, *Olig2*, *Ngn2*, *Lhx3*, *Isl1* and *Chat*. (D) Inferred changes in gene expression over pseudotime from 9000 shortest paths connecting start and end cells (stars in B). Each shortest path was resampledtoalengthof41pseudo-timepointstoenablestatistical measurements of gene expression. Cell IDs color coded according to cell states in A. Quantification of the global rate of change in gene expression identifies three metastable states (light gray) separated by transition states during which the rate of change in gene expression is increased (dark gray). Transition phases are defined as intervals along the pseudo-temporal timeline at which the second derivative of the global gene variation is negative, while metastable states are characterized by a positive second derivative. (E) Gene expression profiles along pseudotime for NP TFs (*Irx3*, *Pax6*, *Nkx6.1* and *Olig2*), gene associated with the transition to MNs (*Ngn2*, *Lhx3* and *Neurod4*) and MN markers (*Isl1/2*, *Tubb3* and *Chat*). (F) Levels of gene expression for *Hes1/5*, *Olig2* and *Ngn2* over pseudotime. Note that *Olig2* expression appears biphasic with upregulation of *Olig2* concommitant to *Ngn2* induction and repression of *Hes1/5* in the transition phase from NP to MN.

Whereas the five cell clusters defined by these modules represented a progressive shift of cell states from early progenitor cells to MNs, the remaining cell cluster exhibited a divergent gene expression signature. In this cluster, many genes contained in Modules 1 and 2 were downregulated but neuronal gene expression was not increased. This cluster exclusively consisted of day 6 cells (Fig 2A). *Nkx2.2* could be detected in some cells of this cluster (Fig S3C), suggesting that it was comprised of cells progressing from a pMN to a more ventral p3 identity. Further differential gene expression analysis on this population identified *Fatty-acid binding protein 7* (*Fabp7*) as enriched in these cells (Fig S3C). Fabp7 levels are markedly upregulated in p3 progenitors *in vitro* and at cervical and brachial levels in embryonic spinal cords at e10.5 (Fig S3D). We therefore conclude that this cluster contains cells progressing from MN to p3 progenitors. Taken together, this suggests our scRNA-seq analysis identifies cells along the MN developmental timeline and partitions these into specific cell types from early NPs to post-mitotic MNs and p3 progenitors.

We next asked whether it was possible to reconstruct the developmental timeline from the transcriptome data. For this we used the 306 genes contained in the five neural gene modules to visualize the developmental trajectory as a pseudo-temporal ordering derived from a consensus of a large number of randomized minimum spanning trees (see Analytical Supplement). The resulting cell graph represents the predicted developmental order of cells based on their transcriptome profile and hence differentiation state (Fig 2B). Strikingly, the five previously characterized cell clusters were ordered on the cell state graphs as expected from the characterization of their gene expression profile (Fig 2C). The graph revealed developmental trajectories originating from *Irx3* expressing early NPs to MN progenitors characterized by *Olig2* expression (Fig 2C). These progenitors then differentiated into MNs via the sequential expression of *Ngn2*, *Lhx3*, *Isl1* and *Chat* (Fig 2C), or into p3 progenitors characterized by *Nkx2.2* and *Fabp7* expression (Fig S3C). To investigate these trajectories in more detail we focused on the developmental trajectory leading from NPs to MNs. To represent changes in gene expression in an unbiased manner, we reconstructed the average gene expression program along pseudotime from the 9000 shortest paths connecting *Irx3* expressing progenitor cells to differentiated MNs on the cell state graph (starred cells in Fig 2B, see Analytical Supplement). Each individual path was resampled to a constant length of 41 pseudotime points (Fig 2D), allowing statistical measurements along the developmental timelines. The outcome was predicted gene expression dynamics during MN differentiation

### Characterization of transcriptional changes during MN differentiation

As a first validation, we asked if the pseudo-temporal ordering reproduced the temporal sequence of well-characterized gene expression changes that lead to MN differentiation. The inferred trajectory correctly predicted the induction sequence of homeodomain and bHLH TFs *Irx3*, *Pax6*, *Nkx6.1*, and *Olig2* involved in ventral patterning of the spinal cord (Fig 2E) (Dessaud et al. 2010; Jeong and McMahon 2005; Chamberlain et al. 2008). Next, we focused on the transition from progenitors to MNs. As expected, this transition was associated with the transient expression of *Ngn2*, *Neurod4* and *Lhx3*, followed by the expression of MN markers including *Isl1/2*, *Tubb3* and *Chat* (Fig 2E). To assess the robustness of these gene expression dynamics, we utilized a bootstrapping approach to ask how dependent these are on individual cells with particular gene expression values (see Analytical Supplement). A total of a 1000 bootstrapped datasets were constructed by randomly drawing cells, with replacement, while maintaining original sample size. Then expression profiles were calculated for each gene in each replicate (Fig S4). To statistically quantify their robustness, we asked how well these profiles were correlated between each pair of replicates (see Analytical Supplement). This analysis revealed a mean Spearman correlation value greater than 0.85 for most genes (Fig S4). This suggests that the observed gene expression dynamics do not depend on the levels of gene expression in specific cells along the pseudo-temporal trajectory and are a robust representation of the gene expression dynamics during MN differentiation.

The process of cell development has been characterised as a series of metastable states defined by a relatively homogenous gene expression program connected by stereotypic transitions (Moris et al. 2016). During these transitions coordinated changes in gene expression occur, often induced in response to a change in signalling. We reasoned metastable states and transition phases should be evident in the pseudo-temporal ordering. Quantifying the variation in gene expression by averaging the normalized derivative of the most dispersed genes’ expression profiles identified these phases (Fig 2D). The three metastable states in which gene expression changes were relatively modest corresponded to early NPs, MN progenitors and MNs. Linking these states were transitions characterized by an increased change in the global gene expression profile. The first transition corresponded to the switch from *Irx3* expressing intermediate progenitors to *Olig2* expressing MN progenitors (Fig 2E), while the second captured the transition of progenitors to post-mitotic neurons (Fig 2E).

We asked whether signatures of signalling pathways driving these transitions could be identified. To this end, we examined the induction and disappearance of canonical target genes for different signalling pathways. As expected, the transition from *Irx3* to *Olig2* coincided with the induction of well-known Shh target genes *Ptch2*, *Hhip1* and *Gli1,* consistent with Shh signalling mediating this transition (Fig S3E). By contrast, the second transition was accompanied by a loss of Notch signalling, marked by the disappearance of *Hes1/5* and induction of markers causing or characteristic of a loss of Notch signalling, including *Numbl*, *Hes6*, *Dll1*, *Ngn2* and *Neurod4* (Fig 2E and Fig S3F). Strikingly, the beginning of this stage coincided with peak expression levels of *Olig2* (Fig 2E,F). This finding raised the possibility that high levels of Olig2 promote neurogenesis, potentially by directly regulating levels of Notch signalling. In summary, the characterization of changes in the transcriptional profile in pseudotime identified distinct metastable cell states and the signalling pathways associated with the transitions between these states.

### *In vitro* and *in vivo* validation of the pseudo-temporal ordering

To extend this approach and validate the predicted timeline we asked whether the data was sufficient to capture fine-grained temporal information that could be tested experimentally. Examination of the transition from *Olig2* expressing progenitors to *Isl1* expressing MNs predicted the transient expression of first *Ngn2*, then *Lhx3* and finally *Isl1* (Fig 2C,E). This is consistent with *in vivo* data indicating that *Lhx3* precedes the expression of other MN markers in the spinal cord (Arber et al. 1999; Tanabe et al. 1998) and a similar sequence of gene expression has been described in an *in vitro* MN differentiation protocol based on embryoid bodies (Rhee et al. 2016; Tan et al. 2016). To confirm this sequence of events *in vitro* we assayed Olig2, Ngn2, Lhx3, and Isl1 on day 6 of differentiation and quantified the levels of expression in individual nuclei (Fig S5A). Comparison of Olig2 and Isl1 levels in individual nuclei revealed a clear trajectory from Olig2-positive, Isl1-negative NPs to Isl1-positive, Olig2-negative MNs. Overlaying the levels of Ngn2 and Lhx3, in the same cells, revealed that both proteins are only transiently expressed along the differentiation trajectory (Fig S5B,C). To confirm the absence of Lhx3 in more mature MNs, we assayed Lhx3, Isl1 and the pan-neuronal marker Tubb3 (Fig S5D). Consistent with the pseudo-temporal ordering, most Tubb3 expressing cells displayed high levels of Isl1 expression but only low levels of Lhx3, while cells with high levels of Lhx3 did not express high levels of Isl1 or Tubb3 (Fig S5D). In summary, these two observations confirm the predictions from the pseudo-temporal ordering and validate the approach for predicting fine-grained changes in the transcriptional program of cells along the differentiation trajectory to MNs.

To further test the reliability of the timeline and demonstrate the validity of the approach for understanding MN differentiation dynamics, we asked if we could predict novel genes involved in MN formation. To this end, we selected genes positively correlated with *Olig2* and *Ngn2* (Fig S5E,F). One gene with a particularly strong relationship was *Zbtb18* (also known as *RP58* or *Zfp238*). Zbtb18 is a zinc-finger TF with a BTB domain. In the brain its loss causes microcephaly and decreased neuronal and increased glial differentiation (Xiang et al. 2011). Less is known about its expression pattern and role in the spinal cord, although *in situ* hybridisation analyses have suggested it is predominantly expressed in ventral progenitors (Oosterveen et al. 2013). As expected, when we assayed Zbtb18 using immunohistochemistry, it was expressed in cells that also expressed Olig2 and Ngn2 (Fig S5G-I). Consistent with this, its expression was detected *in vivo* in the pMN domain at e9.5 in cells that also expressed high levels of Olig2 and Ngn2 (Fig S5J). At e10.5, it was still predominantly expressed ventrally, although no longer confined to the pMN domain (Fig S5K). In summary, this expression pattern further validates the computationally reconstructed MN differentiation timeline.

### Olig2 expression increases as cells commit to MN differentiation

The MN differentiation timeline indicated that *Olig2* expression was induced as *Irx3* was repressed (Fig 2E), consistent with the cross repressive interactions between these two genes (Novitch et al. 2001; Mizuguchi et al. 2001; Chen et al. 2011). This transition demarcated the transition from the first to the second phase identified in the MN timeline. It was noticeable that the expression of *Olig2* appeared biphasic with a marked increase in levels of *Olig2*, which coincided with the transition from the second to the third phase. Moreover, this transition corresponded to the induction of *Ngn2*. This predicted that *Olig2* levels peak at the onset of differentiation before being downregulated as MN identity is elaborated (Fig 2E). To test this prediction, we first examined the levels of Olig2 and Ngn2 in the neural tube of e9.5 and e10.5 embryos, during the period of MN production (Fig 3A-D’’). Consistent with previous studies, we found that at both stages a proportion of Olig2 expressing cells also expressed Ngn2, while a much lower proportion of cells expressed Ngn2 outside the pMN domain (Mizuguchi et al. 2001; Scardigli et al. 2001; Novitch et al. 2001). To test if the levels of Olig2 expression varied in the way predicted by the scRNA-seq data, we quantified levels of Olig2, Ngn2 and the MN marker Mnx1 in nuclei of the pMN domain (Fig 3E-H). This revealed a striking correlation between Olig2 and Ngn2 protein levels in individual cells throughout the pMN domain (Fig 3E,G). Moreover, cells expressing high levels of Olig2 and Ngn2 were differentiating into MNs as measured by the induction of Mnx1 (Fig 3H). This quantification also indicated that Olig2 protein persisted longer than Ngn2 in MNs, as cells co-expressing high levels of Olig2 and Mnx1, but not Ngn2, were observed (Fig 3H). Taken together, these data suggest that high levels of Olig2 correspond to the induction of Ngn2 and the onset of neurogenesis within the pMN domain.

**Fig 3.**
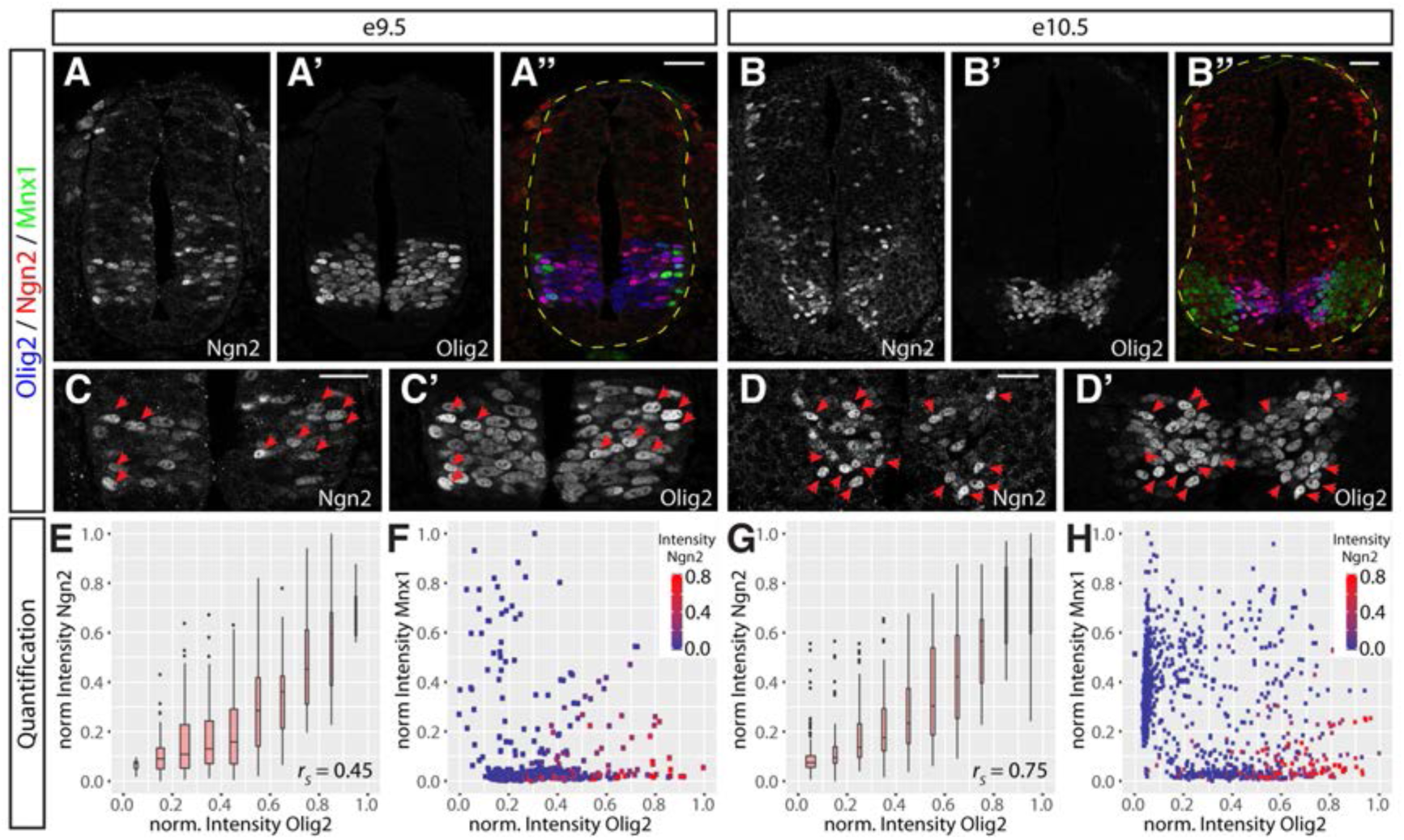
Olig2 expression is higher in Ngn2 expressing progenitors in the pMN domain. (A-B’’) Staining for Ngn2 (A, B), Olig2 (A’, B’) merged with Mnx1 (green in A’’, B’’) in spinal cords at e9.5 (A-A’’) and e10.5 (B-B"). (C-D’) Higher magnification images of the spinal cords shown in (A-B"). Red arrowheads indicate nuclei with elevated levels of Ngn2 and Olig2. (E, G) Positive correlation between Olig2 and Ngn2 protein levels in individual nuclei at e9.5 (E, n = 464 nuclei) and e10.5 (G, n = 1078 nuclei). (F, H) Levels of Olig2, Mnx1 and Ngn2 in individual nuclei throughout the pMN domain at e9.5 (F) and e10.5 (H). Plotting Olig2 versus Mnx1 protein levels reveals a clear differentiation trajectory from Olig2-positive pMN cells to Mnx1-positive MNs. Note that high levels of Ngn2 are only observed in cells with high levels of Olig2 expression. In addition Olig2 protein perdures much longer in Mnx1 positive MNs than Ngn2. Scale bars = 50 μm.

These data prompted us to test directly whether progenitors that expressed high levels of Olig2 were committed to MN differentiation. Since endogenous Olig2 protein disappears rapidly from differentiated MNs, we took advantage of an ESC line in which we fused the fluorescent protein mKate2 to the C-terminus of endogenous Olig2 via a self-cleaving peptide (Fig 4A) (Shcherbo et al. 2009; Szymczak et al. 2004). In these cells, the expression of mKate2 provides a readout of Olig2 levels but the increased stability of fluorescent protein offers a way to mark the progeny of Olig2 expressing cells and estimate Olig2 levels in the progenitor. Control ESC differentiations indicated that Olig2 expression dynamics, protein levels and MN formation were similar in cells containing the engineered or wild-type *Olig2* allele (Fig 4B and Fig S6A-C). Quantification of the mKate2 and Olig2 protein levels in individual nuclei revealed a positive correlation in most cells (Fig 4C-C’’,G and Fig S6D-F). However, we noted a cohort of cells with much higher levels of mKate2 relative to Olig2. Assaying Isl1/2 expression revealed that these cells were MNs (Fig 4D-D’’). Consistent with this, high levels of Isl1/2 and Tubb3 expression were only detected in cells with high levels of mKate2 (Fig 4H and Fig S6A-C). Moreover, mKate2 levels negatively correlated with levels of the NP marker Sox1 (Fig 4E-E’’,I). Thus, MNs indeed progress through a distinct Olig2^HIGH^ state as they exit from the NP state.

**Fig 4.**
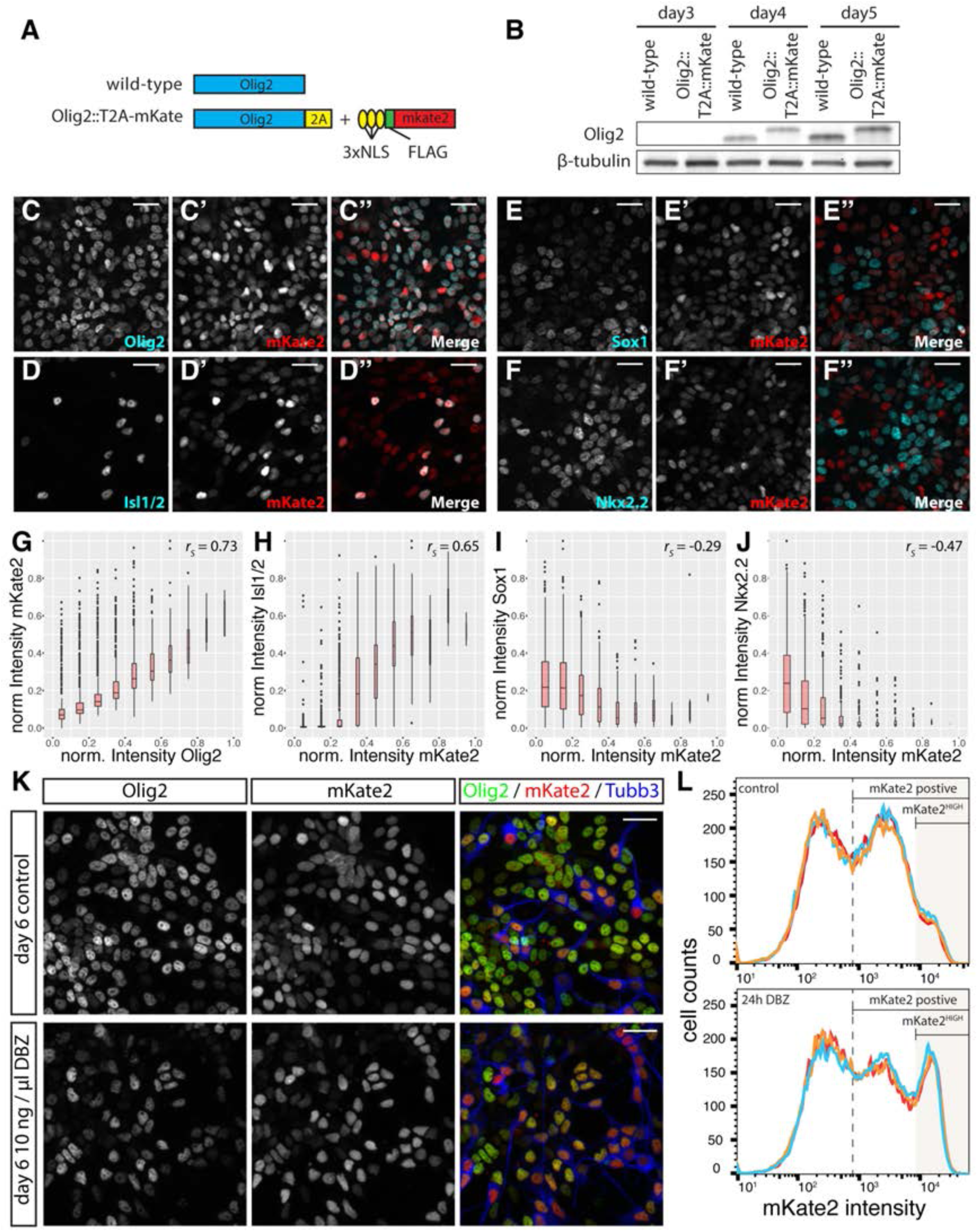
Quantification of a fluorescent Olig2 reporter reveals a marked upregulation of Olig2 prior to MN differentiation. (A) Design of the Olig2-mKate2 reporter. A 3xNLS-FLAG-mKate2 reporter was fused to the C-terminus of endogenous Olig2 via a T2A self-cleaving peptide. (B) Western Blot analysis reveals that the targeted allele shows the same expression dynamics and levels as endogenous Olig2. The targeted allele runs at slightly increased molecular weight due to addition of the T2A peptide (see A). Note that both alleles are targeted in this cell line and consequently no protein of wild-type size was detected. (C-F’’) Immunofluorescence for mKate2 with Olig2 (C-C’’), Isl1/2 (D-D’’), Sox1 (E-E’’) and Nkx2.2 (F-F’’) at day 6 of the differentiations. (G-J) Quantification of protein levels of mKate2 and Olig2 (G, n = 2851 nuclei), Isl1/2 (H, same dataset as G), Sox1 (I, n = 2049 nuclei) and Nkx2.2 (J, n = 2034 nuclei) in individual nuclei. Note the positive correlation between mKate2 and Olig2 and Isl1/2, and negative correlation between mKate2 and Sox1 and Nkx2.2. (K) Inhibition of Notch signaling using 10 ng/µl DBZ causes an increase of neurogenesis. Immunofluorescent staining for Olig2, mKate2 and Tubb3 in control or after 24 hours DBZ treatment at day 6 of the differentiation. (L) Frequency plots of mKate2 fluorescence intensity obtained by flow cytometry reveal a strong increase in the number of mKate2^HIGH^ cells after 24 hours DBZ treatment. Scale bars = 25 µm.

To address whether the transient upregulation of Olig2 expression was specific for the transition from pMN cells to MNs, we quantified levels of mKate2 in Nkx2.2-expressing p3 progenitors (Fig 4F-F’’). During development these progenitors transit through an Olig2-expressing pMN intermediate state before losing Olig2 expression and inducing Nkx2.2 (Chamberlain et al. 2008; Dessaud et al. 2010, 2007). In contrast to the positive correlation between Isl1/2 and mKate2 (Fig 4H), cells expressing high levels of Nkx2.2 had low or undetectable levels of mKate2 expression (Fig 4J and Fig S6F). Thus, distinct Olig2 expression dynamics underlie the progression of pMN cells to MNs and p3 progenitors.

### Inhibiting Notch signaling increases Olig2 expression

These observations raise the question of what upregulates Olig2 prior to MN formation. The Notch signalling pathway is implicated in controlling the rate of neurogenesis and inhibition of Notch signalling in NPs is well known to trigger neuronal differentiation (Artavanis-Tsakonas 1999; Selkoe and Kopan 2003; Shimojo et al. 2011; Louvi and Artavanis-Tsakonas 2006). Furthermore, the inferred MN differentiation trajectory indicated that two canonical effectors of the Notch pathway, Hes1 and Hes5, decreased as cells switched from the early phase of Olig2^LOW^ expression to Olig2^HIGH^. We therefore tested whether inhibiting Notch signalling upregulated Olig2. As expected, inhibition of Notch signalling, through the addition of the β-secretase inhibitor Dibenzazepine (DBZ), for 24 hours between day 5 and day 6 of differentiation caused a substantial increase in the number of neurons observed (Fig 4K, compare Fig S6C and I). Quantifying mKate2 levels using flow cytometry revealed a similarly substantial increase in the number of cells expressing high levels of mKate2 (Fig 4L and Fig S6G). Furthermore, co-staining these cells with the pan-neuronal marker Tubb3 revealed that most of the mKate2^HIGH^ cells were neurons (Fig S6I). To test whether the increase in mKate2 fluorescence is due to increased Olig2 expression upon Notch inhibition, we quantified mRNA levels of *Olig2* and other progenitor and neuronal markers using RT-qPCR after 0, 12 and 24 hours of Notch inhibition (Fig S6J-L). In contrast to other progenitor markers (*Hes1/5*, *Sox2*, *Pax6*), which decreased upon Notch inhibition (Fig S6J), *Olig2* levels peaked at 12 hours before decreasing after 24 hours, (Fig S6K). The observed *Olig2* expression dynamics are strikingly similar to those of other genes previously implicated in MN formation, including *Ngn2* and *Pou3f2* (Fig S6K). Consistent with the increase in the expression of neurogenic markers after 12 hours, we also observed an increase in the expression of neuronal genes 12 hours and 24 hours after Notch inhibition (Fig S6L). Taken together, these data suggest that Notch signalling controls the transition between the distinct phases of *Olig2* expression by restraining *Olig2* expression in MN progenitors.

### Olig2 represses the expression of *Hes1/Hes5*

To test whether the upregulation of Olig2 coincided with the downregulation of Hes1 and Hes5 *in vivo* we examined the expression of these proteins in mouse embryos. Hes1 is broadly expressed by dorsal progenitors that express the homeodomain protein Pax3, as well as floor plate and p3 cells, marked by the expression of Foxa2 and Nkx2.2, respectively (Fig 5B,C,F,G) (Hatakeyama et al. 2004). By contrast, Hes5 is expressed by cells in the intermediate spinal cord, marked by the expression of Irx3 and Pax6 (Fig 5B,D,E). Olig2 expression was first detectable at e8.5, a time at which Hes5 was broadly expressed throughout the ventral neural tube (Fig 5M). Shortly thereafter, Olig2 and Hes5 showed a high degree of co-expression, which coincided with an increase in the number of Olig2 expressing MN progenitors (Fig 5N-R). However, coexpression of Olig2 and Hes5 appeared to be transient, as by e9.5, Hes5 was downregulated in most Olig2+ cells, and few co-expressing cells could be found by e10.5 (Fig 5Q). During this time, Hes1 expression was low or absent in most MN progenitors (Fig 5H-L). The progressive decrease in Hes5 expression from Olig2+ cells was mirrored by a reciprocal increase in Ngn2 expression and, subsequently, the exit of these cells from the cell cycle and the onset of MN differentiation marker expression (Fig5 M’-Q’, Novitch et al, 2001; Miziguchi et al. 2001, Lee et al. 2005). Thus, the transient coexpression of Olig2 and Hes5 *in vivo* marks the pMN state, while the clearance of Hes5 from Olig2-positive cells coincided with the onset of Ngn2 expression and MN differentiation.

**Fig 5.**
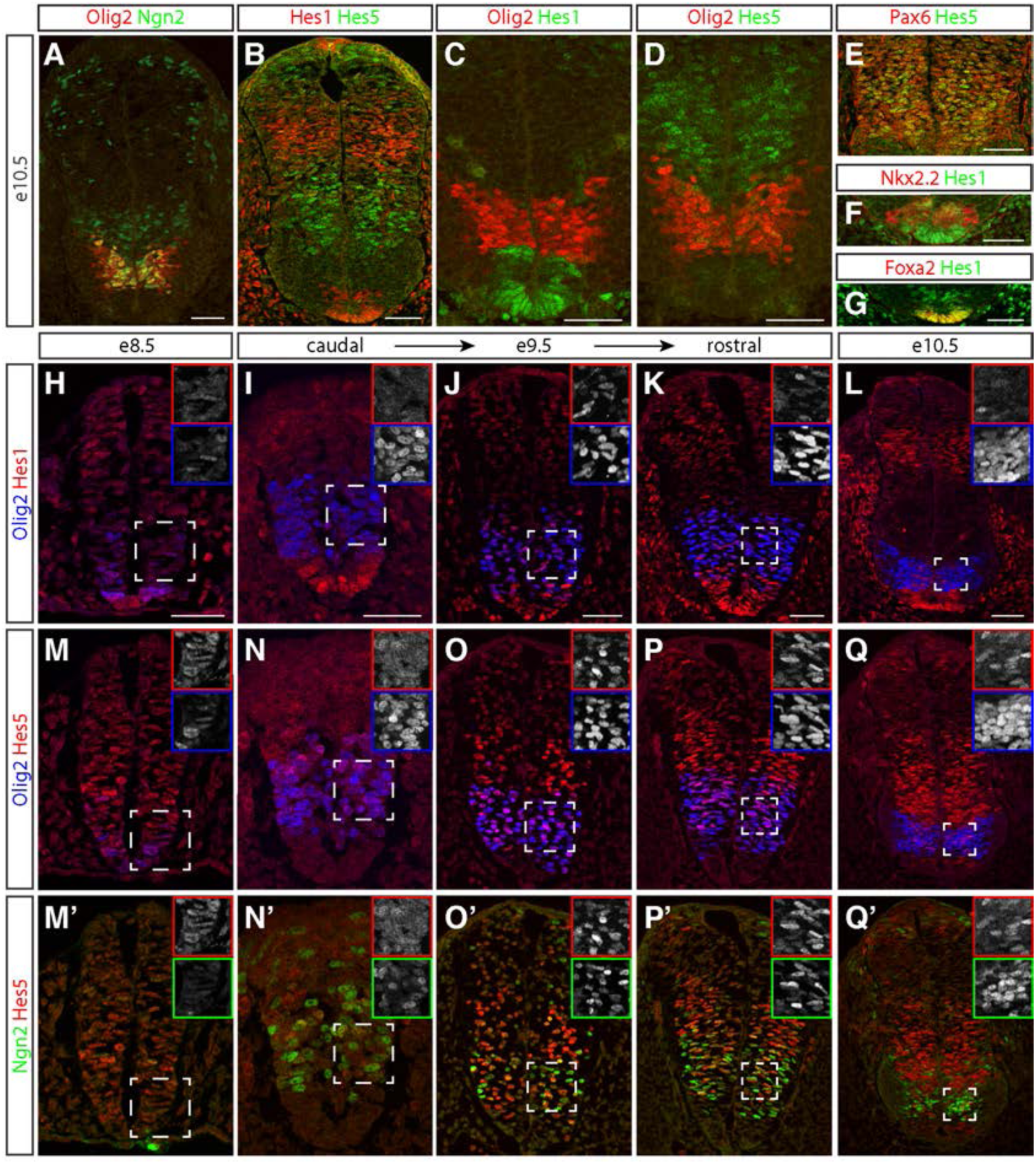
Olig2 and Hes are dynamically expressed in the mouse neural tube. (A-D) Expression patterns of Ngn2 (green in A), Olig2 (red in A,C,D), Hes1 (red in B, green in C) and Hes5 (green in B,D) in the neural tube at e10.5. Note the low expression levels of Hes1/5 and high expression levels of Ngn2 in the pMN domain (compare A,B). (E) Hes5 (green) expression coincides with the expression of high levels of Pax6 (red) in the intermediate neural tube. (F,G) Hes1 expression (green) is readily detected in both Nkx2.2^+^ p3 progenitors (red in F) and floor plate cells labelled by Foxa2 expression (red in G) (H-Q’) Time course of Olig2 (blue), Hes1 (red), Hes5 (red), and Ngn2 (green) expression in neural tubes between e8.5 and e10.5. Multiple panels shown for e9.5 reflect developmental progression from caudal to rostral postions along the neuraxis. Hes1 expression appears to recede from the ventral neural tube upon the onset of Olig2 expression at e8.5 (H) and is thereafter absent from most Olig2+ cells (I-L). Olig2 and Hes5 are initially coexpressed (M,N). Over time, Hes5 expression progressively disappears from the pMN domain (N-Q), and Ngn2 concomitantly increases (N’-Q’). Insets show single channel images of the outlined area for the respective markers. Scale bars = 50 µm.

We next asked whether Olig2 might be responsible for the repression of *Hes1* and *Hes5* using *Olig2*^*Cre*^ knock-in mice (Dessaud et al 2007; Kong et al. 2015). In these mice, the Olig2 coding sequence has been replaced with Cre recombinase (Dessaud et al 2007). Thus, Cre protein expression demarcates the presumptive pMN domain in heterozygous control and in homozygous *Olig2*^*Cre/Cre*^ mutant embryos, which entirely lack Olig2 activity (Fig 6A,E). In controls, the pMN domain was flanked dorsally and ventrally by Hes5 and Hes1 expression, respectively, with little overlap of Cre with either protein (Fig 6C-D’,I). By contrast, *Olig2* mutant spinal cords displayed a marked dorsal expansion of Hes1 and ventral expansion of Hes5 into the pMN domain, such that their expression domains appeared to contact one another (Fig 6G-I). This juxtaposition was associated with a substantial decrease in the number of cells expressing Ngn2 in the pMN domain (Fig 6B,F,I). Thus, Olig2 is required to maintain the boundaries of Hes1 and Hes5 and allow Ngn2 to accumulate within MN progenitors (Fig 6I).

**Fig 6.**
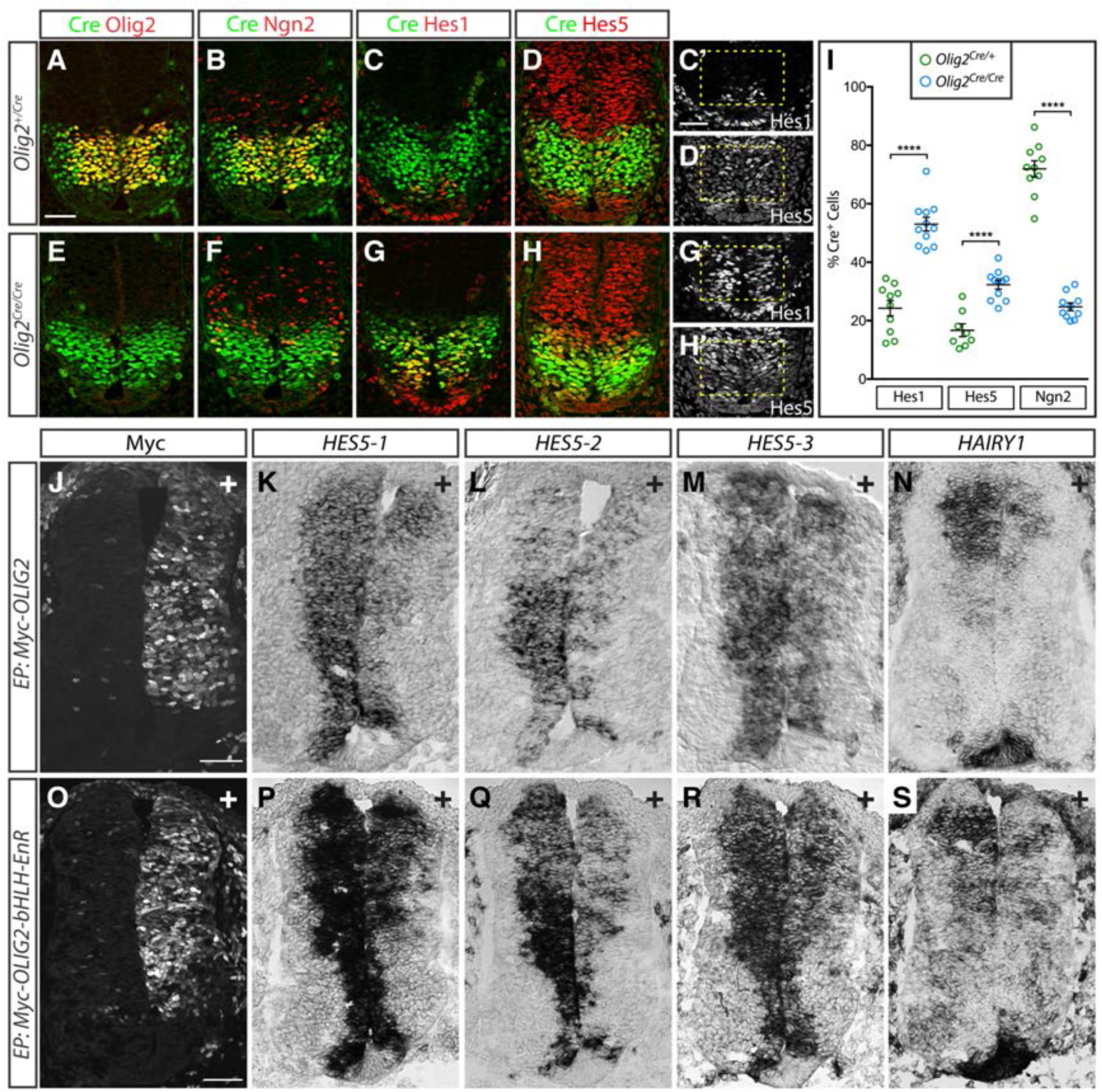
Repression of Hes1/5 in the pMN domain depends on Olig2 activity. (A-D) Expression of Cre (green in A-D), Olig2 (red in A), Ngn2 (red in B), Hes1 (red in C, grey in C’) and Hes5 (red in D, grey in D’) in e10.5 *Olig2*^*Cre*^ heterozygous embryos. (E-H) *In Olig2*^*Cre/Cre*^ homozygous mutants, Hes1 expands dorsally (G, G’) and Hes5 ventrally (H, H’) into the pMN domain, marked by Cre expressed from the *Olig2* locus. The expansion of Hes1/5 coincides with a loss of the high levels of Ngn2 normally seen in the pMN domain. (I) Quantification of Hes1, Hes5 and Ngn2 expression in *Olig2*^*Cre*^ heterozygous and homozygous embryos. The overlap between Cre and Hes1/5 significantly increases in *Olig2*^*Cre*^ homozygotes while overlap between Ngn2 and Cre is strongly reduced. Plot shows the mean ± SEM from multiple sections collected from 3 to 5 embryos for each group. Each section is represented by a single dot with n = 8-11 for each group. **** p < 0.0001, unpaired t-test. (J-S) Electroporation of myc-tagged OLIG2 and an OLIG2-bHLH-Engrailed repressor domain fusion protein in chick neural tubes represses expression of the Hes5 homologues *HES5-1* to *HES5-3* (K-M; Q-R) and the Hes1 homologue *HAIRY1* (N, S). ‘+’ indicates transfected side of the spinal cords. Results are representative of > 5 successfully transfected embryos collected from two or more experiments. Scale bars = 50 µm.

To address whether Olig2 expression was sufficient to repress *Hes1* and *Hes5*, we used *in ovo* electroporation to deliver retroviral expression constructs driving the expression of a myc-tagged form of OLIG2 into the developing spinal cord of Hamburger-Hamilton (HH) stage 11-13 chick embryos. These conditions have been previously shown to increase *NGN2* expression (Novitch et al. 2001). Whereas mammals have a single *Hes5* gene, birds contain three *Hes5* paralogs, termed *HES5-1*, *HES5-2*, and *HES5-3* (Fior and Henrique 2005), clustered at a common genomic locus (Fig 7A). When Olig2 was misexpressed, all three chick *HES5* genes were substantially reduced, as was the chick *Hes1*-related gene *HAIRY1* (Fig 6J-N). Similar results were achieved with misexpression of a dominant repressor form of OLIG2 containing its bHLH DNA binding domain fused to a heterologous Engrailed transcriptional repression domain (Fig 6O-S; Novitch et al. 2001). Based on these results we conclude that Olig2 expression suffices to repress *Hes* gene expression in NPs.

**Fig 7.**
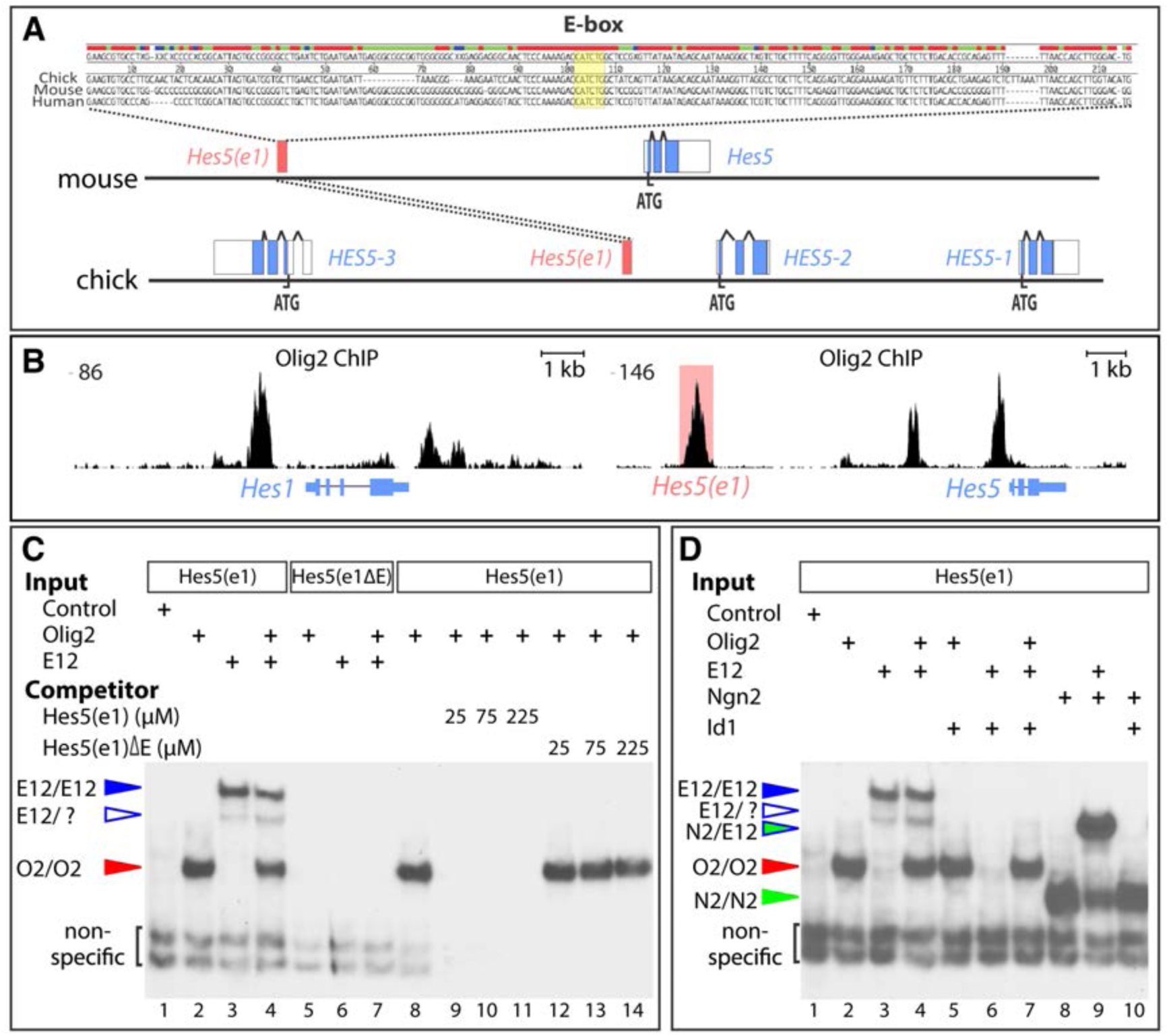
Olig2 binds to an evolutionary conserved element near Hes5. (A) Identification of an evolutionary conserved element containing an E-box in the vicinity of the *Hes5* genomic locus in chick, mouse and human (Hes5(e1)). (B) Analysis of Olig2 Chip-Seq data from Kutejova et al. 2016 reveals Olig2 binding sites in the vicinity of the *Hes1* and *Hes5* genes. The peak corresponding to the Hes5(e1) element is highlighted in red. (C) Electrophoretic mobility shift assays show that both Olig2 and E12 homodimers can individually bind to the Hes5(e1) E-box, and do not form any heterodimeric complexes (lanes 1-4). Positions of the different protein complexes are indicated by colored arrows. Binding depends on the E-box as both proteins fail to bind probes containing an E-box mutation (Hes5(e1ΔE) (lanes 5-7). Olig2 binding to Hes5(e1) can be abolished by the addition of unlabelled Hes5(e1) probes, but not those containing the E-box mutation (lanes 8-14). (D) Id1 inhibits binding of E12, but not of Olig2 or Ngn2, to the Hes5(e1) element. Olig2, E12 and Ngn2 alone or Ngn2/E12 heterodimers can bind the Hes5(e1) element. Mixing Olig2 or Ngn2 with Id1 does not inhibit their homodimeric binding activities (lanes 2, 5, 8, and 10). In contrast, Id1 strongly inhibits binding of both E12/E12 and Ngn2/E12 complexes (lanes 6 and 10). The addition of E12 without and with Id1 does not affect Olig2 binding efficiency (lanes 2, 4, and 7).

Hes proteins and pro-neural bHLH TFs such as Ngn2 act antagonistically in multiple developmental contexts (Kageyama et al. 2007; Shimojo et al. 2011) and ectopic Olig2 expression has been shown to promote Ngn2 expression (Novitch et al. 2001). Two sequences of events could explain the repression of *Hes* genes and induction of Ngn2 in MN progenitors. Either Olig2 represses *Hes1/5* and thereby indirectly induces Ngn2 or, alternatively, Olig2 induces Ngn2, which then antagonizes the expression of the Notch effectors. To distinguish between these possibilities, we investigated if Olig2 represses *Hes5* in *Ngn2* null mutants (Fig S7A-F’’). To this end, we utilized a *Ngn2 knock-in GFP* (*Ngn2*^*KIGFP*^) mouse line, in which an IRES-GFP construct has been inserted into the coding sequence of *Ngn2* (Seibt et al. 2003). Assays at e9.5 and e10.25 revealed that Olig2 expression was maintained (Fig S7A’-F’) and Hes5 repressed in MN progenitors lacking Ngn2 (Fig S7A’’-F’’). Consistent with the observed repression of *Hes5*, GFP expression from the endogenous *Ngn2* locus was strongly elevated in MN progenitors (Fig S7C-F). We therefore conclude that Olig2, not Ngn2, is the main repressor of *Hes* genes in MN progenitors.

These findings raise the question of how important the Olig2-mediated repression of *Hes* genes is for the pattern of neurogenesis in the ventral spinal cord. To address this, we investigated the consequences of preventing Olig2-mediated repression by ectopically expressing chick *HES5-2*, an ortholog of murine *Hes5*, in the ventral spinal cord of chick embryos. Ectopic *HES5-2* expression did not have a noticeable effect on the levels of the progenitor markers SOX2, NKX6.1, PAX6 and OLIG2 (Fig S7G-J). By contrast, and consistent with the well-known anti-neurogenic role of Hes proteins (Ohtsuka et al. 1999; Hatakeyama et al. 2004; Imayoshi et al. 2013), ectopic *HES5-2* resulted in the down-regulation of the pro-neuronal TFs NGN2 and NEUROD4 and of the pan-neuronal marker NEUN (Fig S7K-Q). Of note, cells that maintained NGN2 and NEUROD4 in these experiments usually contained little if any GFP, marking transfected cells, suggesting they were not electroporated (Fig S7N,O). Consistent with this, only a minor fraction of GFP-electroporated cells left the ventricular zone and activated NEUN expression (compare Fig S7M and P). These results suggest that the repression of *Hes* genes is the key mechanism by which Olig2 promotes neurogenesis, and that the anti-neurogenic function of Hes proteins needs to be overcome before *Ngn2* can be induced and neurogenesis initiated. Together, these data indicate that Olig2 plays a critical role repressing the expression of *Hes* genes within MN progenitors to promote the expression of pro-neurogenic bHLH TFs such as Ngn2 and Neurod4 and thereby increases the rate of neuronal differentiation in MN progenitors.

### Olig2 acts directly on a Hes5 regulatory element

The striking effects of Olig2 on *Hes1* and *Hes5* expression prompted us to ask whether Olig2 might directly regulate these genes. Examination of chromatin immunoprecipitation data from mouse NPs revealed several prominent binding sites of Olig2 in the vicinity of the two loci (Fig 7B) (Kutejova et al., 2016, http://www.ebi.ac.uk/ena/data/view/ERX628418). Furthermore, some of these binding sites are in close proximity to previously mapped binding sites for the Notch signalling co-factor RBPJ (Li et al. 2012). Bound regions included sites close to the transcription start sites of the genes and in putative distal regulatory elements (Fig 7B). Aligning genomic sequences of the *Hes5* locus from chick, mouse, and human, indicated that one of the binding sites for Olig2 and RBPJ coincided with a highly conserved ~200 base pair element, hereafter termed Hes5(e1), that is 80% identical between mouse and human, and 53% identical between chick and mouse (Fig 7A,B). This element is 7.9 kilobases (kb) 5’ to the transcriptional start site for *Hes5* in mouse, 10.5 kb 5’ to the transcriptional start site in human, and in the middle of the *HES5* gene cluster in chick.

Like many bHLH proteins, Olig2 binds to canonical E-box DNA response elements with the palindromic sequence CANNTG (Lee et al. 2005). This motif was found within the most conserved central region of the Hes5(e1) element (87% identity between chick and mouse over 46 bp; 98% identity between mouse and human) (Fig 7A). To confirm that Olig2 could bind to the Hes5(e1) element, we performed *in vitro* binding experiments using a probe comprising the conserved central region. *In vitro* translated Olig2 readily bound to the Hes5(e1) E-box, as did other bHLH proteins such as E12 and Ngn2 (Fig 7C,D). These binding activities were abolished when the conserved E-box sequence was mutated (Fig 7C). To test if Olig2 binding activity is enhanced by the presence of E proteins, we mixed Olig2 protein with E12, but found no evidence of either an Olig2:E12:DNA complex or enhanced binding affinity to the Hes5(e1) E-box (Fig 7C). In addition, mixing Olig2 with Id1, a potent competitor for E protein binding, did not diminish Olig2 binding although mixing E12 and Ngn2 with Id1 completely abolished both E12 and Ngn2/E12 binding activities (Fig 7D). The binding of both Olig2 and Ngn2 to Hes5(e1) was further confirmed through chromatin immunoprecipitation experiments (Fig S8A). Taken together, these data indicate that Olig2 homodimers bind directly to a highly conserved Hes5(e1) regulatory element through a single E-box site that may be targeted by other bHLH proteins.

### The Hes5(e1) Element Restricts Gene Expression from the pMN

The observation that Olig2 could bind to the conserved element within the *Hes5* locus prompted us to test whether this element restricted gene expression selectively from the pMN domain. To test this, we generated reporter constructs consisting of Hes5(e1) with or without an intact E-box, upstream of a β-globin minimal promoter driving expression of a nuclear *Enhanced Green Fluorescent Protein (EGFP)* gene (Hes5(e1)-βG::*nEGFP*) (Fig 8C). We co-electroporated these constructs into the chick spinal cord together with a nuclear β-galactosidase (βgal) encoding plasmid (Fig 8A-E). CAG-driven βgal expression appeared to be uniform throughout the dorsalventral axis of the neural tube (Fig 8A,D). By contrast, Hes5(e1)-βG::*nEGFP* activity was spatially restricted, with high levels of expression in the intermediate portions of the neural tube but little if any expression in both ventral and dorsal regions (Fig 8B). Strikingly, the ventral limit of Hes5(E1)-βG::*nEGFP* expression coincided with the dorsal border of the Olig2 expression domain (Fig 8B,F).

**Fig 8.**
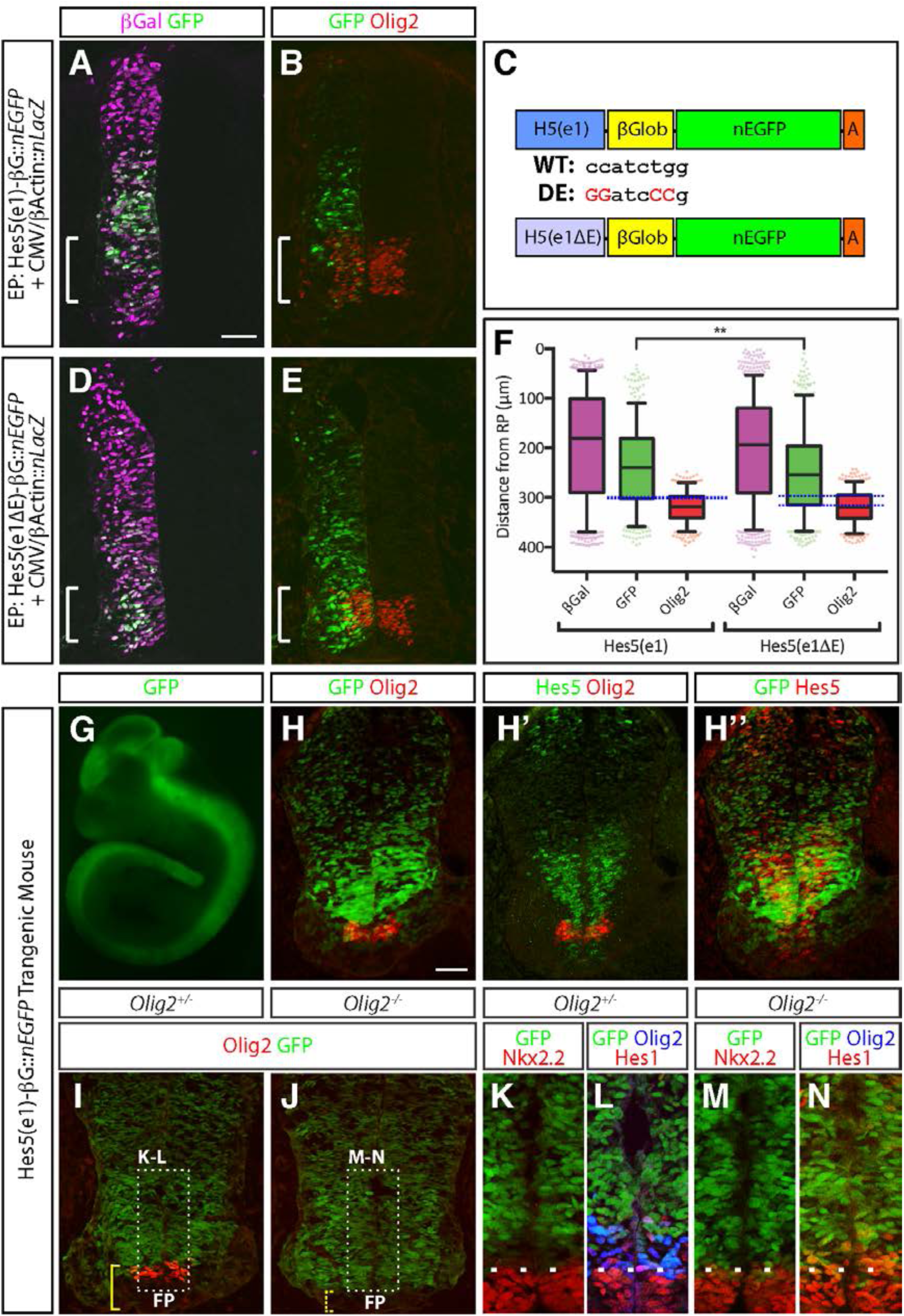
The Hes5(e1) element is required for repression of reporter genes in the pMN domain. (A,B) Co-electroporation of CMV/β-actin::nLacZ and Hes5(e1) reporter plasmids into chick spinal cord. Although electroporation (revealed by β-Gal staining, magenta in A) is uniform along the dorsal-ventral axis, expression of the EGFP-reporter is confined to intermediate parts of neural tube (A,B) and little coexpression of Olig2 and EGFP was detected (B). (C) Design of Hes5(e1) and Hes5(e1ΔE) reporters. The Hes5(e1) element was cloned in front of β-globin minimal promoter to drive *EGFP* reporter gene expression. To test the importance of the E-box in the Hes5(e1) element, critical base pairs for Olig2 binding were mutated (red). (D,E) Co-electroporation of CMV/β-actin::-nLacZ and Hes5(e1ΔE) reporter plasmids into chick spinal cord. In contrast to the Hes5(e1) reporter plasmid, significant coexpression of Olig2 and GFP in the pMN domain is detected (E). Note that E box mutation reduced the basal activity of the reporter, such that longer exposure times were needed to achieve the signals levels seen in the intermediate spinal cord with the nonmutated Hes5(e1) reporter (Fig S8B,C). (F) Scatter dot plots display the dorsal-ventral positions (distance from the roof plate) of individual cells expressing the Hes5(e1) and Hes5(e1ΔE) reporters relative to CMV/β-actin::-nLacZ and Olig2. Results are aggregated from five representative sections taken from five well-electroporated and stage-matched spinal cords. The Hes5(e1ΔE) reporter exhibits a significant ventral shift in its activity and considerable overlap with Olig2 expression (blue dotted box). Lines and error bars indicate mean and interquartile ranges, respectively. *** p = 0.0005, Mann-Whitney test; ns, not significant, p = 0.6649. (G) EGFP-expression in Hes5(e1)-nEGFP whole mount embryos at e10.5. (H-H’’) Cryosections of Hes5(e1)-nEGFP embryos at e10.5 assayed for GFP, Olig2 and Hes5. EGFP expression colocalizes with Hes5 expression (H’’), but not with Olig2 (H). (I-P) Hes5(e1)-nEGFP expression in *Olig2* heterozygous (I,K,L) and homozygous mutants (J,M,N). In *Olig2* heterozygotes, little nEGFP expression can be detected in the Olig2 expression domain, resulting in a pronounced gap between the expression domains of EGFP, Nkx2.2 and Hes1 (K,L). By contrast, the EGFP, Nkx2.2 and Hes1 expression domains directly abut each other in *Olig2* homozygous mutants (M,N).

To determine whether the E-box within Hes5(e1) was essential for this spatially restricted expression pattern, we compared the activity of a Hes5(e1)-βG::*nEGFP* reporter construct in which the E-box had been mutated (Hes5(e1ΔE)-βG::*nEGFP*) (Fig 8D-F). Loss of the E-box substantially reduced the overall activity of the nEGFP reporter compared to the original construct (Fig S8B,C). In addition, nEGFP expression now showed abundant overlap with Olig2 in the ventral spinal cord (Fig 8E,F). Together, these results indicate that the Hes5(e1) element integrates both positive and negative regulatory information through its E-box.

Finally, to test whether Olig2 is responsible for the restriction of Hes5(e1)-βG::*nEGFP* from the pMN domain we generated transgenic mice containing this construct that displayed activity throughout the neuraxis (Fig 8G). In agreement with the chick electroporation data, Hes5(e1)-βG::*nEGFP* activity was spatially restricted, with high levels of expression seen only in intermediate regions of the spinal cord where high levels of Hes5 were expressed (Fig 8H-J). The ventral extent of Hes5(e1)-βG::*nEGFP* activity coincided with the dorsal border of the pMN domain with little overlap between Olig2 and GFP (Fig 8H). By contrast, in *Olig2* mutant embryos the expression of Hes5(e1)-βG::*nEGFP* extended ventrally to reach the dorsal boundary of Nkx2.2, a result that was not seen in control embryos (Fig 8K-P). Together, these data provide evidence that Olig2 represses expression of Hes5 in the pMN domain at least in part through direct interactions with the E-box site within Hes5(e1).

## DISCUSSION

Here, we provide a detailed molecular description of somatic MN differentiation. Single cell transcriptomics defines distinct phases of differentiation and reveals the regulatory relationships that drive progression from NPs to post-mitotic MNs. Experimental validation confirmed these predictions and demonstrated that Olig2 plays a pivotal role coordinating growth and patterning by integrating differentiation and fate determination signals (Fig 9).

**Fig 9.**
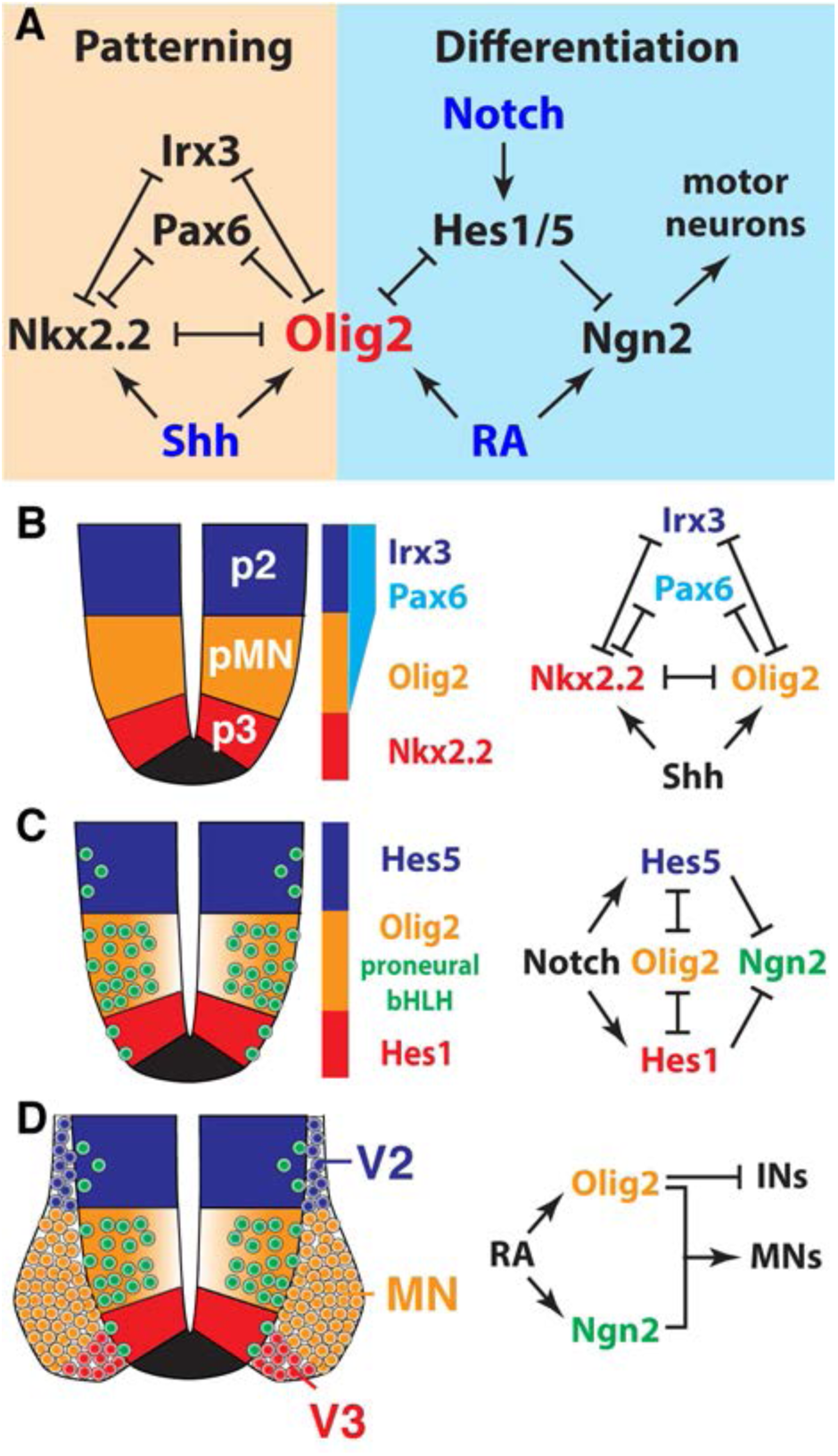
Olig2 coordinates patterning and neuronal differentiation. (A) Proposed model of the Olig2-controlled gene regulatory network. Olig2 does not only act as central organizer for dorsal-ventral patterning in the spinal cord, but also controls the rate of MN differentiation through direct repression of Hes TFs. This leads to a higher levels of Ngn2 expression and consequently a higher rate of neuronal differentiation in the pMN domain compared to adjacent progenitor domains. (B) Olig2 is a core component of the Shh-controlled gene regulatory network that patterns the ventral spinal cord (Balaskas et al. 2012; Cohen et al. 2014). (C) Olig2-mediated downregulation of the Notch effectors Hes1/5 relieves repression of Ngn2 in the pMN domain. (D) Consolidated activities of Ngn2 and Olig2 cause differentiation of NPs to MNs. Olig2 promotes differentiation of MNs through repression of alternative interneuron (IN) cell fates.

### The trajectory of neural progenitor to MN differentiation

Single cell mRNA sequencing is emerging as a powerful tool to reconstruct transcriptional changes in cells during tissue development (Shin et al. 2015; Setty et al. 2016; Trapnell et al. 2014; Treutlein et al. 2016). Here, we use pseudo-temporal ordering of cells based on their expression profile to obtain a high-resolution map of the developmental trajectory of MN differentiation (Fig 2). Examining gene expression along this timeline highlighted the dynamics of signalling pathways and transcriptional networks as cells transit from proliferative progenitors to postmitotic neurons. This computationally reconstructed trajectory accurately recapitulated the known changes in gene expression associated with MN generation *in vivo* and identified features of the dynamics not previously evident. This provides evidence that, by exploiting the inherent heterogeneity and asynchrony of differentiating cells that confound population based assays, scRNA-seq allows the inference of transcriptional dynamics during developmental cell state transitions with high resolution. Moreover, the data illustrate how scRNA-seq analysis of a defined developmental process *in vitro* accurately predicts gene regulatory interactions and transcriptional dynamics *in vivo*.

Examination of the timeline revealed periods of relatively stable gene expression. Punctuating these were transition phases with marked differences in gene expression profiles, which coincided with changes in the signalling status within the cells. This is consistent with the saltatory view of cell fate specification in which differentiation proceeds through a series of metastable states separated by coordinated signal-driven changes in gene expression (Moris et al. 2016). Based on this observation, we used the global rate of change in gene expression in the pseudo-temporal orderings to develop a principled approach that objectively defines these phases. These distinct phases identified in MN differentiation corresponded to known cell types. Expression of *Irx3* marked early, uncommitted NPs, normally located in the intermediate spinal cord. These progenitors transition to pMN cells in response to Shh and retinoid signalling and this was identifiable by the upregulation of *Olig2* and downregulation of *Irx3*. In addition, distinct phases in the acquisition of postmitotic MN identity could be identified with cells expressing markers such as *Lhx3*, *Isl1/2* and *Chat* correctly positioned in the pseudo-temporal ordering.

The temporal ordering provided much greater resolution of the sequence of events leading to MN commitment than previously available. In particular, the transition from MN progenitor to MN was associated with a series of distinct and transient expression changes. This included the induction of well-known pro-neurogenic factors such as *Ngn2*, *Neurod1*, *Neurod4* and *Hes6* (Fig 2E and Fig S3F). Increased expression of *Olig2* was also associated with this stage (Fig 2F). Consequently, the level of *Olig2* expression distinguished two sequential stages in MN progenitors during their differentiation. In the earlier phase, initiated as *Irx3* is downregulated, pMN cells express low or moderate levels of *Olig2*. This is followed by the second phase in which the levels of *Olig2* substantially increase and *Ngn2* becomes expressed at high levels (Fig 2C). *In vivo* analysis, together with the short-term lineage tracing afforded by the Olig2-mKate2 reporter, confirmed that Olig2 upregulation coincided with the commitment to differentiate into postmitotic MNs. By contrast, in the earlier phase of pMN development, the lower levels of Olig2 appeared compatible with the transition of cells to *Nkx2.2 and Fabp7* expressing p3 progenitors (Fig S3C,D). Together, these data provide new insight into the process of MN specification, identifying a series of distinct phases in NP differentiation to fate commitment and highlighting the changes in gene expression that characterise phase transitions.

### Olig2 as a coordinator of neurogenesis

Previous studies have shown that both Olig2 and Ngn2 are required for the elaboration of MN identity and that Olig2 activity induces Ngn2 expression (Novitch et al. 2001; Mizuguchi et al. 2001; Takebayashi et al. 2002; Lu et al. 2000). Consistent with these observations, progenitors in the Olig2 expression domain differentiate at a much higher rate than cells in other progenitor domains of the neural tube (Kicheva et al. 2014; Novitch et al. 2001). Our results suggest a mechanism for this enhanced rate of neuronal differentiation. The canonical Notch effectors Hes1 and Hes5 act to suppress neurogenesis by inhibiting the expression of neurogenic bHLH proteins, thus maintaining NPs in an undifferentiated state (Ohtsuka et al. 1999; Shimojo et al. 2011). Olig2 activity represses *Hes1* and *Hes5* thereby allowing expression of the proneural gene *Ngn2* and downstream effectors such as *Neurod4* (Fig 6 and Fig S7). The ability of Olig2 to repress *Hes5* appears to be direct, as Olig2 binds to a conserved regulatory element within the *Hes5* locus that restricts gene expression from MN progenitors (Fig 7 and Fig 8). Similarly, Olig2 binding sites are found in putative regulatory elements associated with the *Hes1* gene, raising the possibility that this regulatory interaction is also direct. Consistent with a role for Olig2 in promoting neuronal differentiation, the levels of Olig2 transcript and protein peak at the onset of neurogenesis, concomitant with the induction of Ngn2 *in vivo* and *in vitro* (Fig 2 and Fig 3). These findings therefore establish a mechanism by which neural patterning and neurogenesis intersect. In this view, by modulating the Notch pathway, the Shh and retinoid-dependent induction of Olig2 not only specifies MN identity but also determines the rate at which these progenitors differentiate, thus imposing the distinctive kinetics of MN production (Fig 9).

This model is surprising as previous studies suggested antagonistic activities for Olig2 and Ngn2 during the induction of neuronal target genes (Lee et al. 2005). Both Olig2 and Ngn2 have been shown to heterodimerise with E47 and bind to E-box elements but with opposing activities (Lee et al. 2004, 2005). In addition, similar to Id proteins, Olig2 proteins could potentially sequester E proteins (E12 and E47) from forming heterodimeric Ngn2/E-protein complexes that activate transcription (Samanta and Kessler 2004; Lee et al. 2005). The sequential expression of Olig2 and Ngn2 has been proposed as a potential mechanism to reconcile the inhibitory activity of Olig2 on neurogenesis with the high rate of neurogenesis in the pMN domain (Lee et al. 2005). However, our results suggest that this mechanism is unlikely to apply to the differentiation of MNs in the spinal cord for several reasons. The higher temporal resolution provided by the pseudo-temporal ordering indicated that primary Ngn2 target genes such as *Dll1* and *Neurod4* are induced when the rate of Olig2 expression is maximal in cells (Fig 2E and Fig S3F). Furthermore, Olig2 protein perdures longer in differentiating MNs than Ngn2, resulting in significant co-expression of Olig2 and early MN markers such as Lhx3 and Mnx1 in Ngn2-negative cells (Fig 3H and Fig S5B,C). Hence, instead of sequential expression of these proteins, these results suggest that Ngn2 is capable of mediating neurogenesis despite the presence of high Olig2 levels.

A potential solution to this puzzle may be that the activities of both Olig2 and Ngn2 are regulated by phosphorylation. Olig2 phosphorylation at specific Ser/Thr residues regulates its choice of dimerization partner, intracellular localization and DNA binding preference for open and closed chromatin (Sun et al. 2011; Meijer et al. 2014; Li et al. 2011; Setoguchi and Kondo 2004). Indeed, homodimeric complexes of Olig2 appear to mediate Hes5 repression (Fig 7C,D). Furthermore, the cell cycle kinases CDK1/2 have been proposed to phosphorylate Olig2 at Ser14, priming Olig2 for further phosphorylation at multiple Ser residues (Zhou et al. 2017). This phosphorylation appears to regulate the preference of Olig2 for open or closed chromatin and, thus, strongly influences its biological activity (Meijer et al. 2014). The declining levels of CDK1/2 during neurogenesis may similarly affect Olig2 activity to overcome its inhibitory role on neuronal gene expression. Similarly, phosphorylation of Ngn2 affects its stability and interaction with Lim-homeodomain TF complexes and E-proteins (Ma et al. 2008; Hindley et al. 2012; Ali et al. 2011; McDowell et al. 2014). Thus, additional post-translational events extend the regulatory interactions between both proteins beyond stochiometric interactions through protein-protein binding and competition for DNA binding sites. Notably, some of the relevant phosphorylations are performed by protein kinase A (PKA) and glycogen-synthase kinase 3 (GSK3), kinases linked to the activity of the Shh pathway, which appears to peak at the initiation of MN differentiation (Balaskas et al. 2012; Li et al. 2011; Ma et al. 2008; Kicheva et al. 2014). Connecting the activity of these neurogenic TFs to the activity of the Shh pathway would allow a tight coupling between MN generation and overall developmental dynamics dictated by the dynamics of morphogen signalling.

The pseudo-temporal ordering indicated that although *Olig2* levels peaked at the onset of MN differentiation, expression then decreased rapidly, prior to MNs reaching the next metastable state along the differentiation trajectory, characterised by the induction of mature MN markers such as *Isl2* and *Chat* (Fig 2). This suggests that *Olig2* may need to be downregulated to allow progression of MN differentiation. Consistent with this, overexpression of Olig2 has been shown to inhibit the generation of MNs and to directly repress genes associated with MN identity, such as *Hb9* (Lee et al. 2004, 2005). Furthermore, the addition of Olig2 to canonical reprogramming factors decreases the efficiency of conversion from fibroblasts to spinal MNs (Son et al. 2011). Thus, Olig2 upregulation can promote MN generation by initiating differentiation but its downregulation is needed to complete the switch from progenitor to post-mitotic neuron. These dynamics of Olig2 expression may help impose directionality to differentiation and ensure the correct temporal sequence of gene expression occurs as MNs mature.

### Oscillation of bHLH TFs in the spinal cord

The maintenance of NPs in the brain has been ascribed to the oscillatory expression of Hes and proneural bHLH TFs (Imayoshi et al. 2013; Shimojo et al. 2008). The Hes proteins are proposed to generate the oscillations by negatively regulating their own expression as well as Ngn2 and Ascl1 (Takebayashi et al. 1994; Imayoshi and Kageyama 2014; Shimojo et al. 2011). This phenomenon results in Hes1 and proneural bHLH TFs exhibiting reciprocal expression phases at an equivalent frequency (Shimojo et al. 2008; Imayoshi et al. 2013). Oscillations in the levels of the Notch ligand Dll1 have been reported in spinal cord progenitors (Shimojo et al. 2016). In cortical progenitors, fluctuations in Olig2 levels have also been documented, but these oscillations occur at a significantly slower frequency (Imayoshi et al. 2013) and may therefore be regulated by a different mechanism.

Although we did not specifically investigate the occurrence of bHLH oscillations in the spinal cord *in vitro* or *in vivo*, our results may shed light on this question. The Hes5(e1) element can be bound by both Olig2-Olig2 repression dimers and Ngn2-E12 activation complexes (Fig 7B,C and Fig S8A). It is notable that mutation of the E-box in this element reduced the overall level of reporter activity in the spinal cord (Fig S8B,C), at the same time as disrupting its spatial restriction from the pMN (Fig 8). These data are consistent with a model in which positive activators, such as Ngn2 or other E-box binding factors, could interact with Hes5(e1) to directly elevate Hes5 expression, which would in turn serve to repress Ngn2 expression, thereby contributing to alternating phases of Hes5 and Ngn2 expression. In this regard, Olig2 binding and repressing Hes5 through this element would interrupt the oscillator, allowing Ngn2 expression to reach its maximal levels and neuronal differentiation to commence. Thus, by inserting itself into the Notch regulated neural differentiation program, Olig2 shuts down Notch activity, ensuring MN development proceeds in a spatially and temporally controlled manner. This reconciles stochastic and oscillatory models of neuronal differentiation with the spatially predetermined pattern of neuron production observed in the spinal cord.

To examine Olig2 expression, we used the relatively long half-life fluorescent protein mKate2 introduced into the *Olig2* genomic locus. Quantification of the levels of mKate2 and Olig2 revealed a striking correlation between both proteins in NPs (Fig S6D-F). This argues against Olig2 oscillations in these cells, as oscillatory behaviour would be expected to decrease the correlation between both proteins. Although further investigation is necessary, the data are consistent with out-of-phase oscillations between Ngn2 and Hes5, while Olig2 levels steadily increase in MN progenitors over time. Understanding these relationships will provide insight into the transition from MN progenitor to differentiation.

### The Notch pathway regulates Olig2 expression and Shh signaling

Besides promoting neurogenesis, inhibition of Notch effectors also appears to be important for dorsal-ventral patterning of the neural tube and the consolidation of pMN identity. Patterning of the ventral neural tube is mediated by a gene regulatory network that interprets both levels and duration of Shh signalling (Balaskas et al. 2012; Cohen et al. 2014; Dessaud et al. 2010; Sagner and Briscoe 2017). Previous studies have suggested that Notch signalling influences patterning of the ventral spinal cord by promoting the activity of the Shh pathway (Kong et al. 2015; Stasiulewicz et al. 2015). Consistent with this, overexpression of *HAIRY2*, the chick homologue of murine *Hes1*, causes a downregulation of *Olig2* and induction of *Nkx2.2* in the pMN domain (Stasiulewicz et al. 2015). Similarly, sustained activation or inhibition of the Notch pathway causes, respectively, a ventral expansion or recession in p3 progenitors, located ventral to the pMN (Kong et al. 2015). Here, we show that besides modulating Shh activity, Notch signalling can also regulate expression levels of Olig2. Conversely, Olig2 represses the canonical Notch effector Hes5 and could thereby negatively regulate the levels of Shh signalling in the pMN domain. Thus, Olig2 may consolidate pMN identity not only by direct repression of other progenitor markers, but also indirectly by modulating levels of Shh signalling through its effect on Notch pathway.

Taken together, our data reveal a tight coupling between the gene regulatory networks that control patterning and differentiation in the ventral spinal cord. This highlights the pivotal role of *Olig2* in this process, which not only acts as central organizer of dorsal-ventral patterning in the spinal cord, but also as developmental pacemaker for MN formation. The Olig2-mediated repression of Notch pathway targets provides a molecular mechanism for the much higher rate of neurogenesis observed in the pMN domain compared to the rest of the spinal cord and thereby explains the spatial and temporal patterns of neurogenesis observed in the neural tube. These findings raise the question of whether similar mechanisms also apply in other progenitor domains in the neural tube.

## EXPERIMENTAL PROCEDURES

### Animal Welfare

Animal experiments in the Briscoe lab were performed under UK Home Office project licenses (PPL80/2528 and PD415DD17) within the conditions of the Animal (Scientific Procedures) Act 1986. Animals were only handled by personal license holders. *Olig2*^*Cre*^ and *Ngn2*^*KIGFP*^ knock-in/knockout mice were used as previously described (Seibt et al. 2003; Dessaud et al. 2007), and interbred to create Olig2 or Ngn2 mutant embryos. All mice in the Novitch lab were maintained and tissue collected in accordance with guidelines set forth by the UCLA Institutional Animal Care and Use Committee. Fertilized chicken eggs were acquired from AA McIntyre Poultry and Fertile Eggs, incubated and electroporated as previously described (Gaber et al. 2013).

### Differentiation of NPs from mouse ESCs

NPs were differentiated as described previously (Gouti et al. 2014). In brief, HM1 (Thermo Scientific), DVI2 and Olig2::T2A-mKate2 ESCs were maintained in ES cell medium with 1000 U/ml LIF on mitotically inactivated mouse embryonic fibroblasts (feeder cells). DVI2 cells were generated by integrating a 8xGBS-H2B::Venus Shh pathway reporter into the HPRT locus of HM1 cells and used for all ESC experiments except 4-color stainings in Fig S5A,B, which rely on HM1 cells, and experiments in Fig 4 and Fig S6 which were conducted using the Olig2::T2A-mKate2 reporter cell line.

For differentiation cells were dissociated in 0.05% Trypsin (Gibco) and replated onto tissue culture plates for 25 minutes to remove feeder cells. Cells staying in the supernatant were spun down and resuspended in N2B27 medium at a concentration of 10^6^ cells / ml. 45000 cells were plated onto 35 mm CellBind dishes (Corning) precoated with 0.1% Gelatine solution in 1.5 ml N2B27 + 10 ng / ml bFGF. At D2 medium was replaced with N2B27 + 10 ng / ml bFGF + 5 µM CHIR99021 (Axon). At D3, and every 24 hours afterwards, medium was replaced with N2B27 + 100 nM RA (Sigma) + 500 nM SAG (Calbiochem). For Notch inhibition differentiations were treated at day 5 with N2B27 + 100 nM RA + 500 nM SAG + 10 ng/µl DBZ (Tocris Biosciences) for 24 hours. Cells were washed with N2B27 medium at later medium changes when many dead cells were detected in the dish.

### qPCR analysis

mRNA was extracted using RNeasy Mini Kit (Qiagen) according to the manufacturer’s instructions. 1.5 - 2 µg of RNA was used for reverse transcription using Super-Script III First-Strand Synthesis kit (Invitrogen) with random hexamers. Platinum SYBR Green qPCR mix (Invitrogen) was used for amplification on a 7900HT Fast Real Time PCR machine (Applied Biosystems). Expression values were normalized against β-actin. Three independent repeats of each RT-qPCR time-course were performed and three independent samples at each time point of each repeat were analyzed. For a complete list of used primers see Table S1. qPCR data presented in Figs 1, S1, and S6 shows one representative repeat and shows mean ± standard deviation. Heatmap in Fig S2B was plotted using Graphpad Prism 7.

### Immunofluorescent stainings

Cells were washed using N2B27 medium and PBS (Gibco) and then fixed in 4% paraformaldehyde in PBS at 4°C for 20 minutes. After fixation cells were washed twice with PBS and stored in a fridge till stainings were perfomed. For staining, cells were washed three times in PBS containing 0.1% Triton X-100 (PBS-T). Primary and secondary antibodies were diluted in PBS-T + 1% BSA. Cells were incubated with primary antibodies overnight at 4°C, then washed three times for 5-10 minutes in PBS-T, incubated with secondary antibodies for 1 hour at room temperature, and washed again three times in PBS-T. Stainings were mounted using ProLong Gold Antifade reagent (Life Technologies). Mouse and chicken spinal cord tissues were fixed with 4% paraformaldehyde, cryoprotected in 30% sucrose, sectioned, and processed for immunohistochemistry or in situ hybridization as previously described (Sasai et al. 2014; Gaber et al. 2013).

Antibodies against a peptide in the C-terminal portion of mouse Hes5 (APAKEPPAPGAAPQPARSSAK, aa 127-147) were raised in rabbits and guinea pigs (Covance). The rabbit serum was affinity purified and used at 1:8000, and the crude guinea pig serum at 1:16000. Additional primary antibodies were used as follows: goat anti-β-galactosidase (Biogenesis 4600-1409 1:2000), mouse anti-Cre (Covance Covance MMS-106P, 1:2000), rabbit anti-Dbx1 (kind gift of Susan Morten and Thomas Jessell, 1:8000), rabbit anti-Fabp7 (Abcam ab32423, 1:2000 or Chemical AB9558, 1:2000), rat anti-FLAG (Stratagene 200474, 1:1500), chicken anti-GFP (Abcam ab13970, 1:20000), sheep anti-GFP (AbD Serotec 4745-1051, 1:800), rabbit anti-Hes1 (Ito et al. 2000, 1:1000), mouse anti-Hoxc6 (Santa Cruz Biotechnology sc-376330, 1:250), mouse anti-Hb9 (DSHB, 1:40), mouse anti-Isl1/2 (DSHB, 1:100), goat anti-Isl1 (R&D AF1837, 1:1000), rabbit anti-Lhx3 (Abcam ab14555, 1:500), mouse anti-NeuN (Rbfox3, Chemicon/Millipore MAB377, 1:1000), rat anti-chick Neurod4 (NeuroM, Bylund et al. 2003), goat anti-Ngn2 (Santa Cruz Biotechnology sc-19233, 1:500), mouse anti-Ngn2 (5C6, Lo et al. 2002, 1:50), guinea pig anti-chick Ngn2 (Skaggs et al. 2011, 1:2000), mouse anti-Nkx2.2 (DSHB, 1:25), mouse anti-Nkx6.1 (DSHB, 1:100), rabbit anti-Olig2 (Millipore AB9610, 1:1000), guinea pig anti-mouse Olig2 (Novitch et al. 2003, 1:20000), guinea pig anti-chick Olig2 (Novitch et al. 2001, 1:8000), rabbit anti-Pax6 (Millipore AB2237, 1:1000), mouse anti-Pax6 (DSHB, 1:25), goat anti-Sox1 (R&D AF3369, 1:500), goat anti-Sox2 (Santa Cruz Biotechnology sc-17320, 1:2000), rabbit anti-Sox2 (Bylund et al. 2003, 1:2500), rabbit anti-TagRFP (Evrogen AB233, 1:1000), rabbit anti-Tubb3 (Covance PRB-435P, 1:2000), mouse anti-Tubb3 (Covance MMS-435P, 1:1000), rabbit anti-Zbtb18 (Proteintech 12714-1-AP, 1:1000).

Secondary antibodies used throughout this study were raised in donkey. Alexa488, Alexa568, Cy3, and Dylight 647-conjugated antibodies (Life Technologies or Jackson Immunoresearch) were diluted 1:1000, Alexa647 conjugated antibodies (Life Technologies) 1:500. Cy5 conjugated antibodies (Jackson Immunoresearch) 1:700, CF405M donkey anti-guinea pig secondary antibody (Sigma) 1:250.

### Image acquisition and analysis

Immunofluorescent images of ESC-derived NPs were acquired using a Zeiss Imager.Z2 microscope equipped with an Apotome.2 structured illumination module and a 20x magnification lens (NA = 0.75). 5 phase images were acquired for structured illumination. For each image z-stacks comprised of 12 sections separated by 1 µm were acquired. Maximum intensity projection was performed in Fiji.

Cryosections were documented using a Leica SP5 confocal microscope equipped with a 40x oil objective, or Zeiss LSM5, LSM700, or LSM800 confocal microscopes and Zeiss Apotome imaging systems equipped with 10x, 20x, and 40x oil objectives. For nuclear staining intensity measurements 3-4 individual sections separated by 1 µm were analysed. Nuclei segmentation and intensity measurement were performed in CellProfiler. Data was normalized and plotted using R. Other images were processed and manually quantified using Fiji and Adobe Photoshop imaging software.

### Single Cell Sequencing

NPs were dissociated using 0.05% Trypsin (Gibco), spun down in ES-medium, resuspended, washed and spun down in 10 ml PBS (Gibco). Afterwards, cells were resuspended in 1ml N2B27 and filtered into a FACS tube (Falcon). The Fluidigm C1 platform was used to capture individual cells using 96 small or medium IFC chip. Cells were diluted in the range of 250 000-400 000 cells per ml for chip loading. Capturing efficiency was evaluated by manually inspecting each capture site on the chip using the automated NanoEntek JuLi cell imager. Only capture sites containing single cells were processed for library preparation and sequencing. Single cell full-length cDNA was generated using the Clontech SMARTer Ultra Low RNA kit on the C1 chip using manufacturer-provided protocol. ArrayControl RNA Spikes (AM1780) were added to the cell lysis mix as recommended in the Fluidigm protocol. Libraries were prepared using the Illumina Nextera XT DNA Sample Preparation kit according to a protocol supplied by Fluidigm, and sequenced on Illumina Hiseq 4000 for 75bp paired-end runs.

### Generation of Olig2::T2A-mKate2 ESC line by CRISPR

pNTKV-T2A-3xNLS-FLAG-mKate2 was generated by cloning a T2A-3XNLS-FLAG-mKate2 cassette into pNTKV using HpaI and HindIII restriction sites. To integrate the T2A-3xNLS-FLAG-mKate2 cassette at the 3’ end of the Olig2 open reading frame, a donor vector comprising app. 2.8 kb upstream and 5 kb downstream of the Stop-Codon was constructed. For CRISPR/Cas9 mediated homologous recombination, a short guide RNA (sgRNA) sequence (CGGCCAGCGGGGGTGCGTCC) was cloned into pX459 (Addgene) according to (Ran et al. 2013).

For electroporation DVI2 ESCs were grown in 2i medium + LIF. 4µg of both plasmids were electroporated into 4*10^6^ cells using Nucleofector II (Amaxa) and mouse ESC Nucleofector kit (Lonza). Afterwards, cells were replated onto 10 cm CellBind plates (Corning) and maintained in 2i medium + LIF. For selection cells were first treated with 1.5 µg/ml Puromycin (Sigma) for two days and afterwards with 50 µg/ml Geniticin (Gibco) until colonies were clearly visible. Individual colonies were picked using a 2 µl pipette, dissociated in 0.25% Trypsin (Gibco) and replated onto feeder cells in ES-medium + 1000 U/ml LIF in a 96 well plate. Correct integration of the T2A-3xNLS-FLAG-mKate2 transgene was verified using long-range PCRs.

### Western Blots

Cells were lysed in RIPA buffer supplemented with protease inhibitors. 10 µg of total protein was loaded per lane. Antibodies used were rabbit anti-Olig2 (Millipore AB9610, 1:3000) and mouse anti-β-tubulin (Sigma T4026, 1:2000). Secondary antibodies were donkey anti-mouse IRDye 800CW and donkey anti-rabbit IRDye 680RD (both Licor). Blots were scanned using an Odyssey Scanner (Licor).

### Flow Cytometry

To quantify the number of Olig2::T2A-mKate2 positive cells, NPs were trypsinized in 0.05% Trypsin (Gibco) and spun down in ES medium. Cells were resuspended in PBS (Gibco) supplemented with the live-cell staining dye Calcein Violet (Life Technologies) according to the manufacturer’s instructions. Flow analysis was performed using a Becton Dickinson LSRII flow cytometer. HM1 or DVI2 cells were differentiated in parallel and analyzed using the same settings to estimate the number of mKate2 positive cells in Fig 4L and Fig S6H. The threshold for counting cells as mKate2 positive was set to the intensity for which 0.5% of cells without the mKate2 transgene were counted as positive. 30000 events were recorded for each replicate. To count cells as mKate2^HIGH^ cells in Fig 4L and Fig S6G, a threshold was set to the shoulder visible in the histograms of the control conditions depicted in Fig 4L. This shoulder typically appears at day 6 and is not present in day 5 cells. Based on the flow cytometry data in Fig S6C and the correlation between mKate2 fluorescence and Isl1/2 expression shown in Fig 4H, we infer that most mKate2^HIGH^ cells are motor neurons. Cells were counted as mKate2^HIGH^ if their mKate2 intensity exceeded this threshold.

For analysis of Tubb3 stainings by flow cytometry (Fig S6A-C, I), cells were fixed for 20 minutes in 2% PFA on ice, washed three times in PBS-T and incubated with Alexa647-conjugated anti-Tubb3 antibody (BD Pharmingen, 1:10) in PBS-T + 1% BSA for one hour at room temperature. Cells were afterwards washed 3x in PBS-T and endogenous mKate2 fluorescence and Tubb3 staining in 10000 cells were quantified by flow cytometry. Data was analyzed using FlowJo.

### Electrophoretic Mobilty Shift assays

pCS2+ plasmid expression vectors for *Olig2*, *E12*, *Ngn2*, and *Id1* (Novitch et al. 2001) were transcribed and translated *in vitro* using the Promega TNT Coupled Wheat Germ Extract System. Programmed extracts were mixed as indicated in a buffer containing 100 mM Hepes pH 7.6, 25 mM KCl, 1.5 mM MgCl2, 0.2 mM EDTA, 2.5% glycerol, and 100 ng poly dIdC and incubated for 15 min at room temperature. ^32^P-dCTP-labeled probes were generated by Klenow end-labeling of double stranded oligonucleotides containing the native mouse Hes5(e1) sequence (forward 5’-ggccgCTCCCAAAAGAC<u>CATCTG</u>GCTCCGTGTTATAA-3’; reverse 5’-actagTTATAACACGGAGC<u>CAGATG</u>GTCTTTTGGGAG-3’) or an E-box mutated version (forward 5’ ggccgCTCCCAAAAGAggATCccGCTCCGTGTTATAA-3’; reverse 5’-actagTTATAACACGGAGCggGATccTCTTTTGGGAG-3’). The E-box sequence is underlined. Lower case indicates substitutions and added flanking sequences. Samples were incubated with labeled probes for 15 min before resolving on a 4.5% polyacrylamide gel and subsequent autoradiography. Probe competition was achieved by incorporating unlabeled oligonucleotide probes in the binding reaction mix.

### Hes5(e1) Transgenic Assays

Mouse and chick *Hes5(e1)* genomic DNA fragments were amplified by PCR using the following primers: mouse-forward 5’-gag<u>gcggccgc</u>CGGTTCCCACACTTTGGT-3’; reverse 5’-gag<u>actagt</u>CACAGTCCCAAGCTGCTTAAA-3’ and chick- forward 5’-gag<u>gcggccgc</u>TGCGTTTCCCATACTTTTCC-3’; reverse 5’-gag<u>actagt</u>TCTGGCCTTGAAGCTAGGAG-3’, lower case indicates added flanking sequences with restriction enzyme sites (NotI and SpeI) underlined. PCR products were digested and cloned into a reporter construct containing the β-globin basal promoter, a nuclear *EGFP* coding sequence, and a bovine growth hormone polyadenylation sequence (βG::nGFP, Lumpkin et al. 2003). Mutations of the *Hes5(e1)* E-box were generated through splicing by overlap extension PCR. Chick embyos were co-electroporated with chick *Hes5(e1)* constructs along with a plasmid vector producing nuclear-tagged β-galactosidase under the control of the cytomegalovirus enhancer and β-actin promoter. Embryos were collected, fixed, and cryosectioned. Sections were stained using antibodies to β-galactosidase and Olig2 and fluorescent secondary antibodies. GFP levels were measured based on its native fluorescence. Images were collected and positions of cells expressing each marker rendered using the spots function of the Bitplane Imaris 8.4 imaging suite. Calculated positions were exported and processed using Microsoft Excel and Graphpad Prism 7 software.

Transgenic mice expressing the mouse Hes5(e1)-βG::*nGFP* reporter were generated with the assistance of the University of Michigan Transgenic Animal Model Core by microinjection of purified plasmid DNA into fertilized eggs obtained by mating (C57BL/6 X SJL)F1 female mice with (C57BL/6 X SJL)F1 male mice, and subsequent transfer to pseudopregnant recipients. Analysis was conducted on both embryos collected from the primary reporter injections and offspring collected from matings of a transgenic line that passed through the germline.

### Electroporation and In situ hybridization

Chick embryos were electroporated at Hamburger-Hamilton (HH) stages 11-13 with RCAS-myc-tagged chick *Olig2* and *Olig2*-bHLH-Engrailed plasmid constructs (Novitch et al., 2001) and collected at HH stages 21-23. Spinal cord sections were hybridized with digoxigenin-UTP labeled RNA probes generated from plasmid DNA or templates generated by PCR. 3’ UTR sequences for chick *HES5-1* and *HES5-3* were amplified from spinal cord cDNA with the following primers: *HES5-1*, forward 5’-GCG<u>GAATTC</u>AGGGAAGCTCTCACTTAGTGAAC-3’ and reverse 5’-GCG<u>CTCGAG</u>ATACCCTCCTGCTGAAGACATTTGC-3’; *HES5-3*, forward 5’-GCG<u>GAATTC</u>GCCAAGAGCACGCTCACCATCACCT-3’ and reverse 5’-GCG<u>CTCGAG</u>CTACACAGCTTGAGTTATGGTTTAG-3’ and directionally cloned into the pBluescript. Underlined sequence indicates restriction enzyme sites incorporated into the primers. Chick *HAIRY1* and *HES5-2* 3’ UTR riboprobes were generated as previously described (Gaber et al. 2013).

### Chromatin Immunopreciptation-PCR

ESC-derived MN progenitors were dounce homogenized and sonicated in 3 ml lysis buffer (1% SDS, 50 mM Tris pH 8.0, 20 mM EDTA, 1 mM PMSF, and 1X Complete protease inhibitor cocktail (Roche). 150 μg of lysate DNA were used per immunoprecipitation reaction and mixed with 3-5 µl of either rabbit anti-Olig2 antibodies (Millipore AB9610), rabbit anti-Ngn2 serum (generous gift of Dr. Soo-Kyung Lee, Oregon Health Sciences University), normal rabbit sera, or purified rabbit IgG. Antibody-chromatin complexes were collected using Dynabeads protein A (Invitrogen), washed and eluted DNA used for RT-qPCR in triplicate using the following primer pairs: *Hes5(e1)* forward 5’-CTGCTTCTGAATGAATGAGGGCGG-3’ and reverse 5’-AGCAGACGAGCCCTTTATTGCTCT-3’; *Hes5(e2)*, a non-conserved element 3’ to the *Hes5* coding exons that contains an E box, forward 5’-AGATGGCTCAGCGGTTAAGAG-3’ and reverse 5’-CCATGTGGTTGCTGGGATTTG-3’. Fold enrichment for each region was calculated as compared to normal rabbit serum or purified IgG.

## AUTHOR CONTRIBUTIONS

AS, ZG, JD, Conception and Design, Acquisition of Data, Analysis and interpretation of data, Drafting or revising the article, Contributed unpublished, essential data or reagents; JK, DR, CP, SW, MM, NMG, Acquisition of Data, Analysis and interpretation of data, Contributed unpublished, essential data or reagents; JB, Conception and Design, Analysis and interpretation of data, Drafting or revising the article; BN, Conception and Design, Acquisition of Data, Analysis and interpretation of data, Drafting or revising the article

## ACKNOWLEDGEMENTS

We are grateful to Leena Bhaw and Abdul Sesay for excellent support with single cell sequencing, Supraja Varadarajan for assistance with Imaris image processing and Vicki Metzis for help with processing and visualizing ChIP-Seq data. We thank David Anderson, Thomas Edlund, Thomas Jessell, Soo-Kyung Lee, Susan Morton, and Tetsuo Sudo for reagents; Deborah Keller and Mina Gouti for providing the DVI2 mouse ESC line; Francois Guillemot for kindly providing *Ngn2*^*KIGFP*^ mice; Lorena Belen Garcia Perez, Teresa Rayon Alonso and Christopher Demers for comments on the manuscript. The Hes5 antisera were generated in Thomas Jessell’s laboratory with support from the Howard Hughes Medical Institute and NINDS. AS has received funding from an EMBO LTF (1438-2013), HFSP LTF (LT000401/2014-L) and the People Programme (Marie Curie Actions) of the European Union’s Seventh Framework Programme FP7-2013 under REA grant agreement n° 624973. Work in JB’s lab was supported by the Francis Crick Institute which receives its funding from Cancer Research UK (FC001051), the UK Medical Research Council (FC001051), and the Wellcome Trust (FC001051; WT098326MA). Work in BN’s lab was supported by the UCLA Broad Stem Cell Research Center, the NINDS (R01NS053976, R01NS072804, and R01NS085227), the March of Dimes Foundation (5-FY06-7), and the Whitehall Foundation (2004-05-90-APL).

## SUPPLEMENTAL FIGURES + LEGENDS

**Fig S1.**
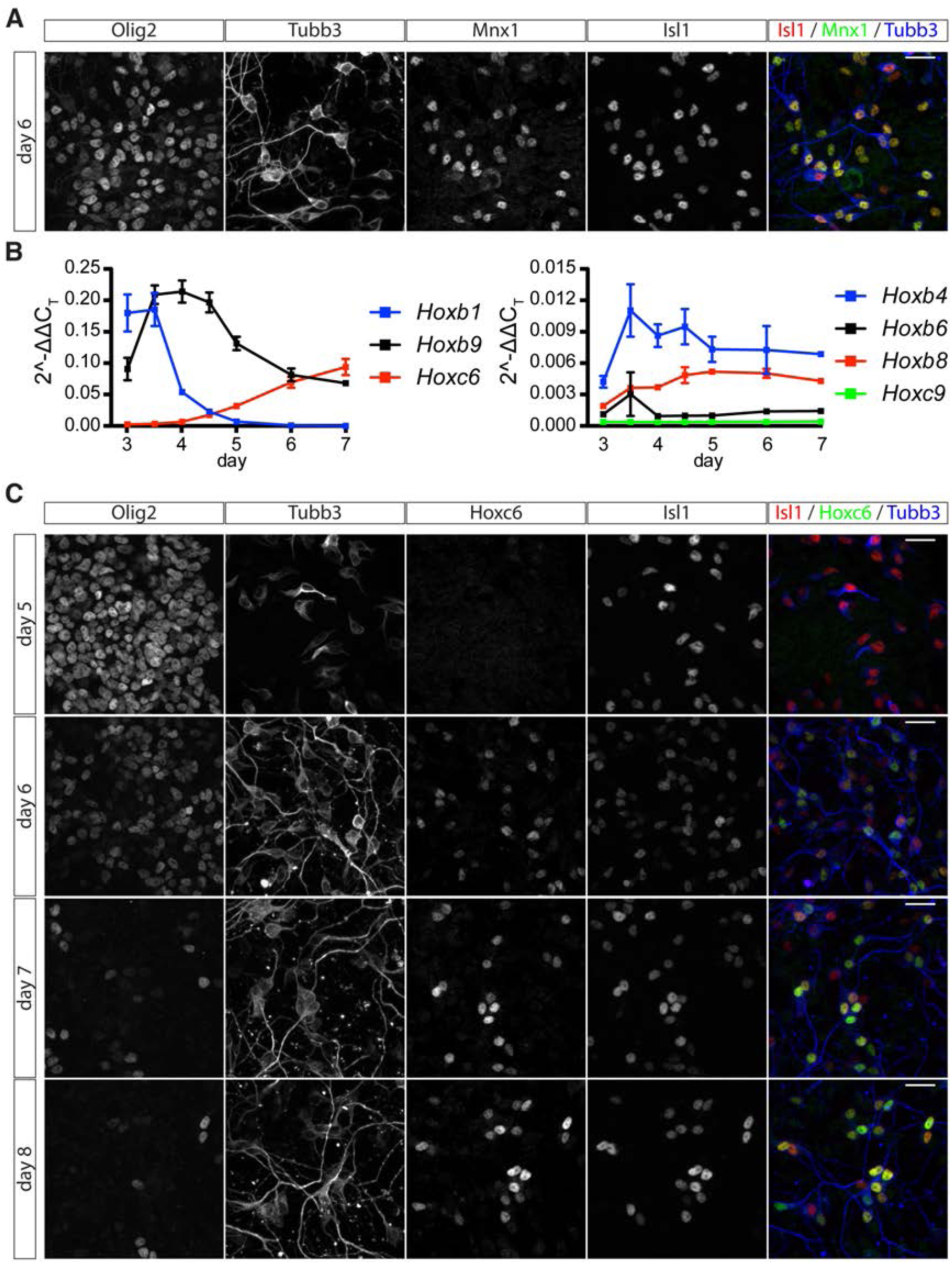
Characterization of Hox gene expression in NPs and MNs. (A) Expression of the somatic MN marker Mnx1 in MNs at day 6. (B) RT-qPCR analysis of *Hox* genes expression levels from day 3 to day 7 (C) Hoxc6 expression in MNs characterized by Isl1 and Tubb3 expression from day 6 to day 8. Scale bars = 40 µm

**Fig S2.**
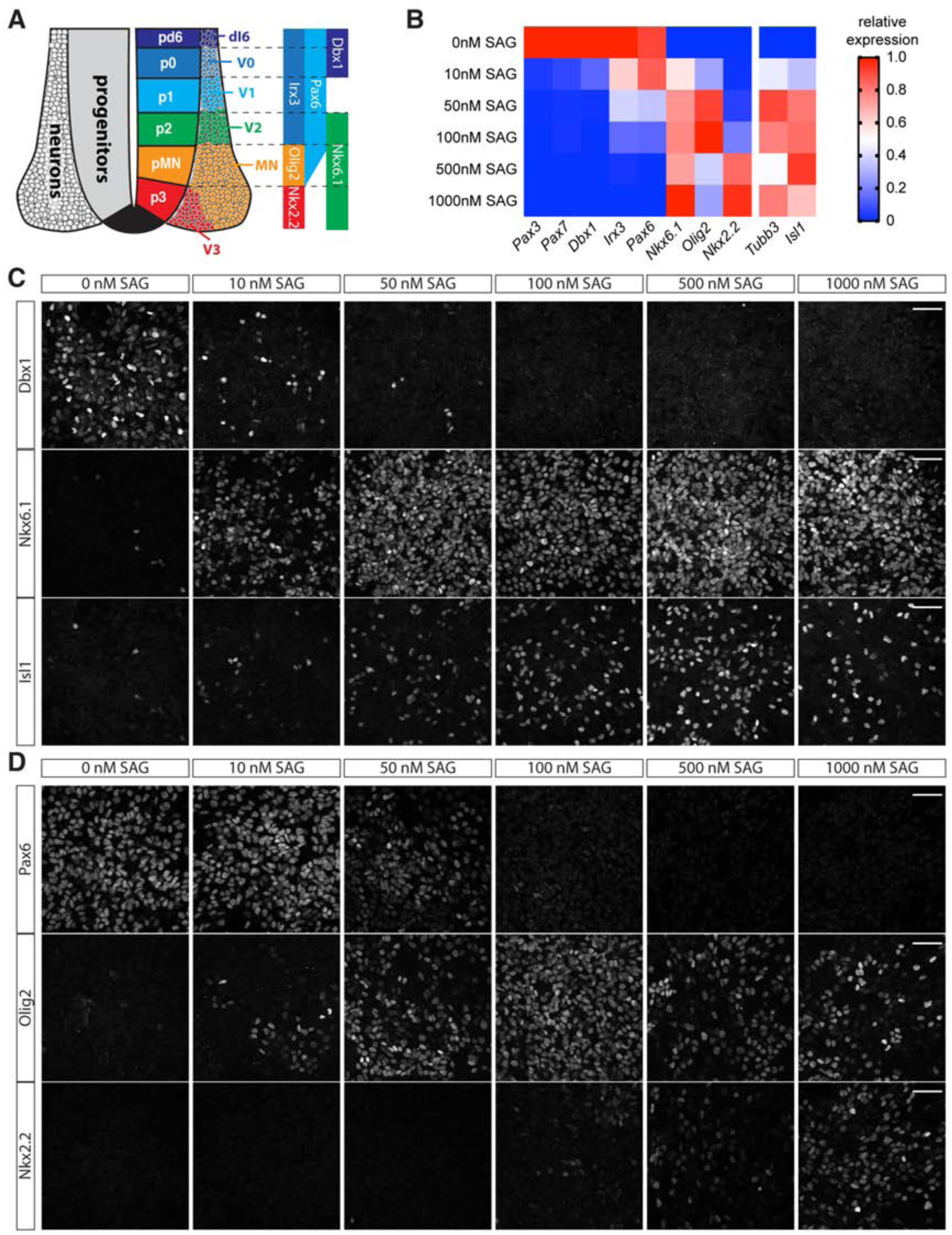
Establishment of different NP identities by different levels of Shh pathway activation. (A) Schematic of the embryonic spinal cord. Expression domains of TFs defining NP domains are indicated. (B) RT-qPCR analysis of day 6 differentiations treated with 0-1000 nM SAG after day 3. (C) Expression of Dbx1, Nkx6.1 and Isl1 in day 6 differentiations treated with the indicated concentrations of SAG (D) Expression of Pax6, Olig2 and Nkx2.2 in day 6 differentiations treated with the indicated concentrations of SAG Scale bars = 40 µm

**Fig S3.**
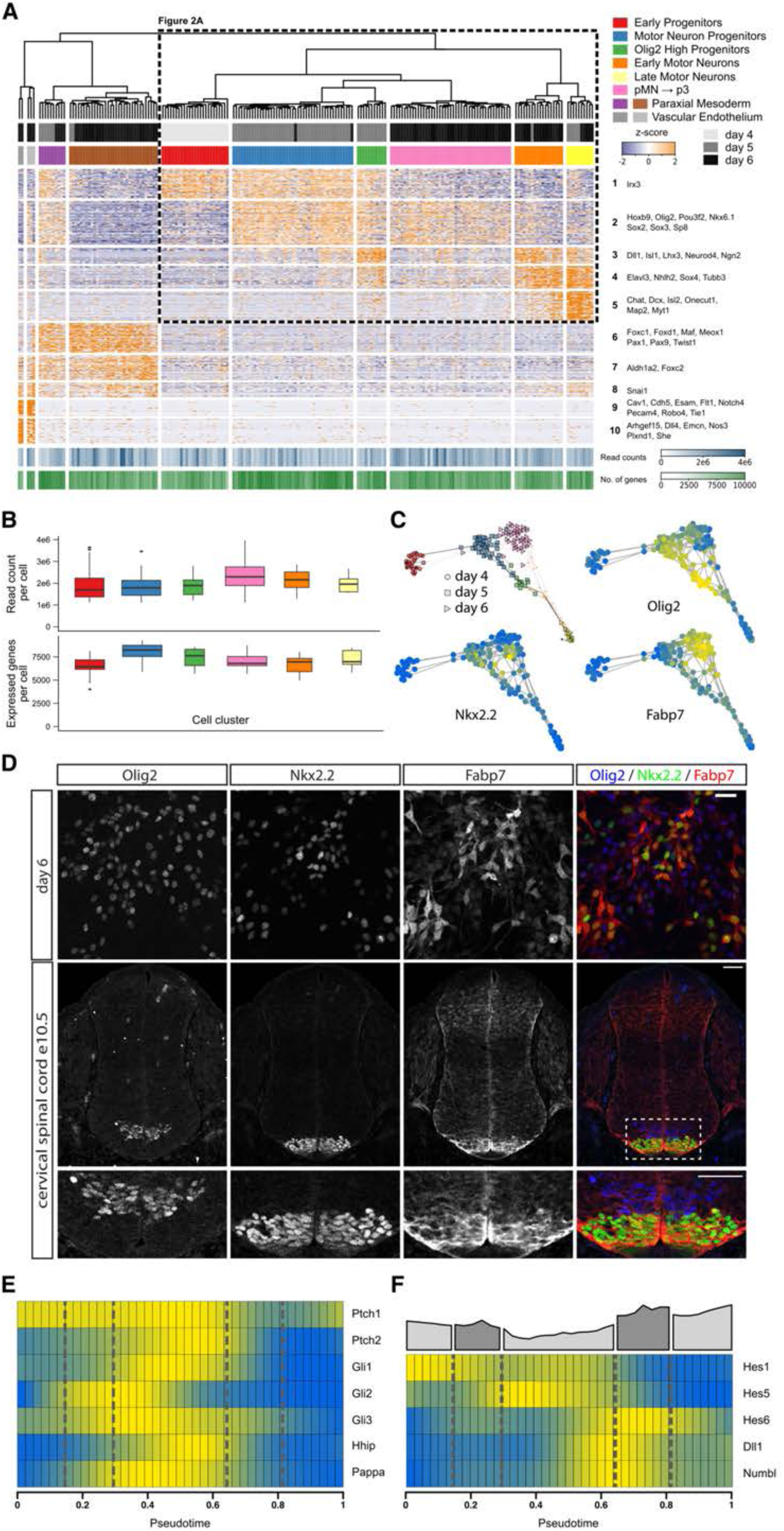
Identification of gene modules and cell states by hierarchical clustering of single cell sequencing data. (A) Identification of cell states by hierarchical clustering from 202 cells based on 10 identified gene modules. Genes characteristic for the individual modules are indicated. Boxed region corresponds to the heatmap in Fig 2A. (B) Quantifications of read counts per cell (top) and number of expressed genes per cell (bottom) for neural cell states identified by hierarchical clustering. Colors of the graphs match cell states in Fig 2A and Fig S3A. (C) Cell state graphs color coded for the expression levels of *Olig2*, *Nkx2.2* and *Fabp7*. (D) Analysis of Olig2, Nkx2.2 and Fabp7 expression in differentiations at day 6 (top row) and e10.5 embryonic spinal cords (bottom row) confirms higher Fabp7 expression levels in p3 progenitors. (E) The transition phase from early *Irx3* NPs to *Olig2* NPs correlates with the induction of Shh target genes *Ptch2*, *Gli1*, *Hhip*. (F) Inhibition of Notch signalling, revealed by decreasing expression levels of *Hes1/5* and expression of the Notch ligand *Dll1* and the pathway inhibitors *Hes6* and *Numbl*, identifies the cell state transition from NPs to MNs. Scale bars = 25 µm (D, top row) and 50 µm (D, bottom row).

**Fig S4.**
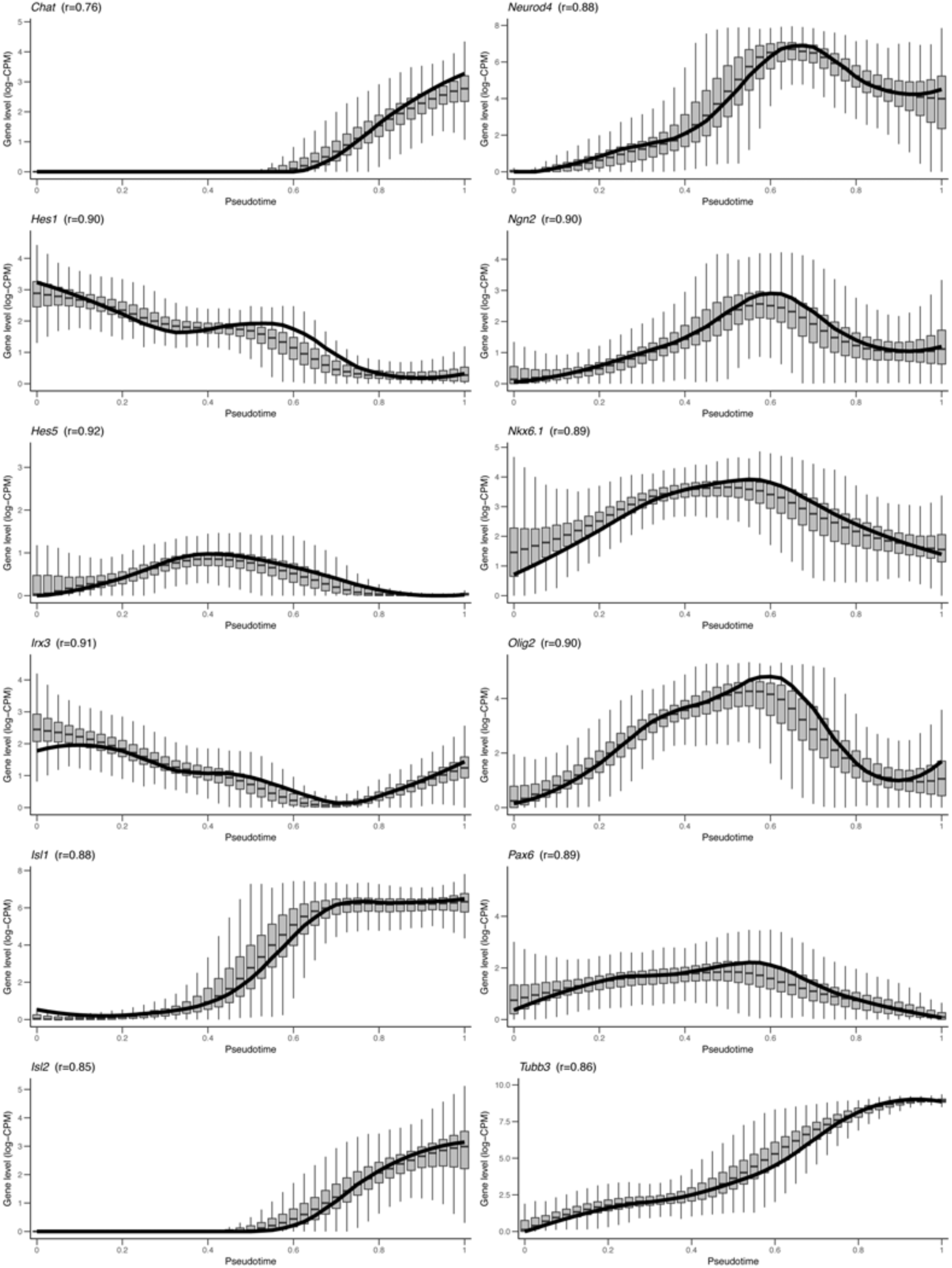
Robustness analysis of gene expression dynamics by bootstrapping. Solid lines indicate original gene levels shown in Fig 2E,F and S3E,F. At each pseudotime point, box plots indicate median, first and third quartiles of the gene level distribution obtained from 1000 bootstrapped datasets. Whiskers indicate the largest and smallest values no further than 1.5 times the interquartile range taken from the hinge. The associated correlation coefficient, r, is the average of the Spearman correlation coefficients over all pairs of boostrap replicates.

**Fig S5.**
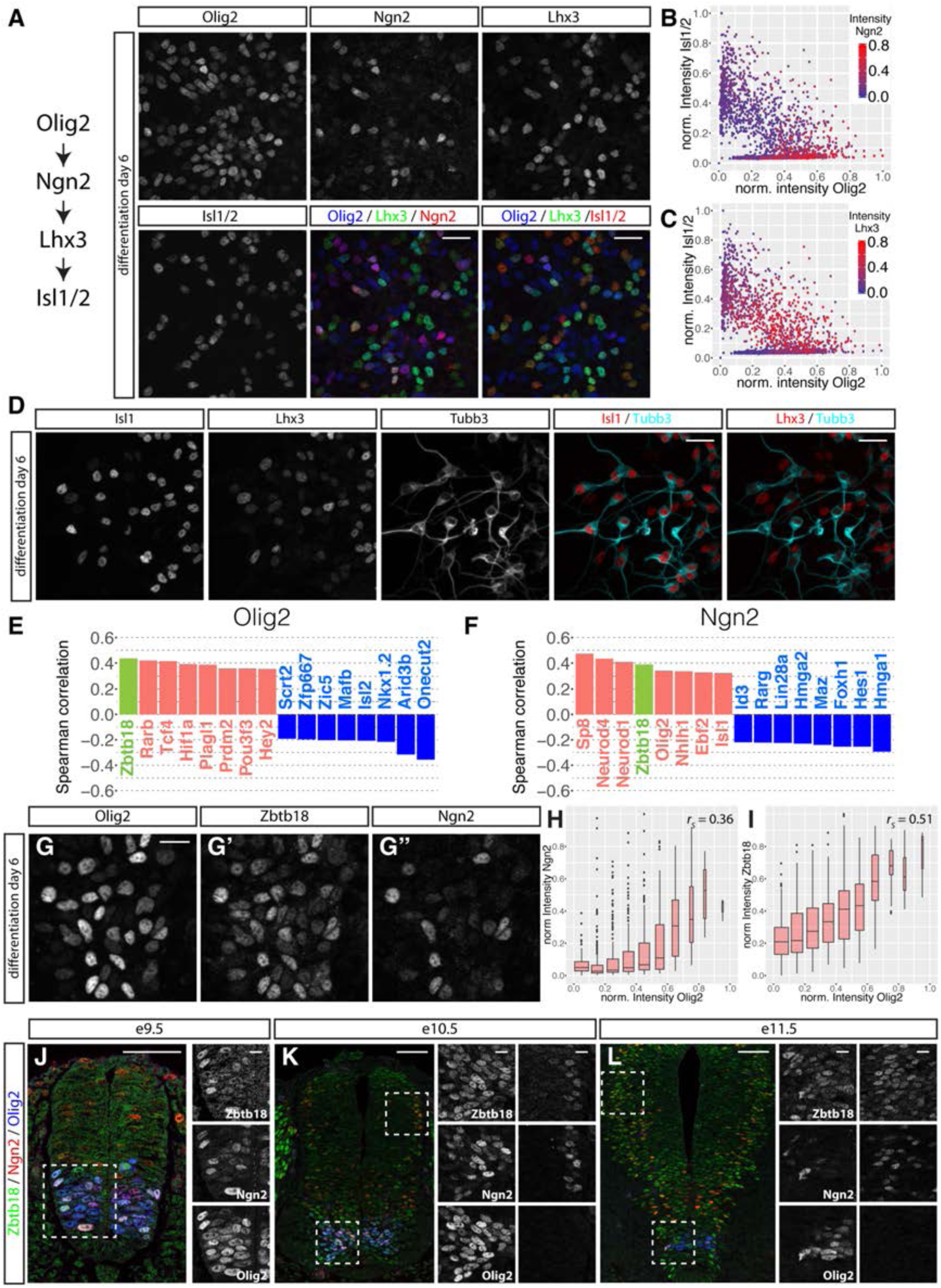
Validation of predictions from the pseudo-temporal ordering. (A) Sequential expression of Olig2, Ngn2, Lhx3 and Isl1 during MN differentiation revealed by immunofluorescent staining for these markers at day 6 of differentiation. (B,C) Quantification of levels of Olig2, Isl1, Ngn2 (color code in B) and Lhx3 (color code in C) reveals a clear differentiation path from Olig2^HIGH^ cells to MNs and sequential induction of Ngn2 and Lhx3 during this process (n = 2236 nuclei). (D) Staining for Isl1, Lhx3 and Tubb3 reveals high levels of Tubb3 expression in Isl1-positive but not Lhx3-positive MNs at day 6 of differentiation. This is consistent with the earlier MN stage of Lhx3 MNs. (E, F) Most positive and negative Spearman-correlated transcription factors for *Olig2* (E) and *Ngn2* (F) reveals *Zbtb18* (green in E-G) as a novel gene involved in MN formation. (G-G’’) Immunofluorescent staining for Olig2 (G), Zbtb18 (G’), and Ngn2 (G’’) at day 6 of differentiation. (H, I) Quantification of levels of Olig2, Ngn2 (H) and Zbtb18 (I) in individual nuclei reveals a good correlation between these markers (n = 1431 nuclei). (J-L) Analysis of Olig2, Ngn2, and Zbtb18 expression in neural tubes at e9.5 (J), e10.5 (K) and e11.5 (L). Note that Ngn2 and Zbtb18 are expressed in cells with high levels of Olig2 at e9.5 and e10.5, but not at e11.5 (left insets in K-M). In addition, Zbtb18 and Ngn2 are co-expressed in nuclei at the edge of the progenitor domain in dorsal areas of the neural tube at e10.5 (L) and e11.5 (M) (right insets). Scale bars = 25 µm in (A,D), 10 µm in (G) and insets in J-L, 50 µm in J-L

**Fig S6.**
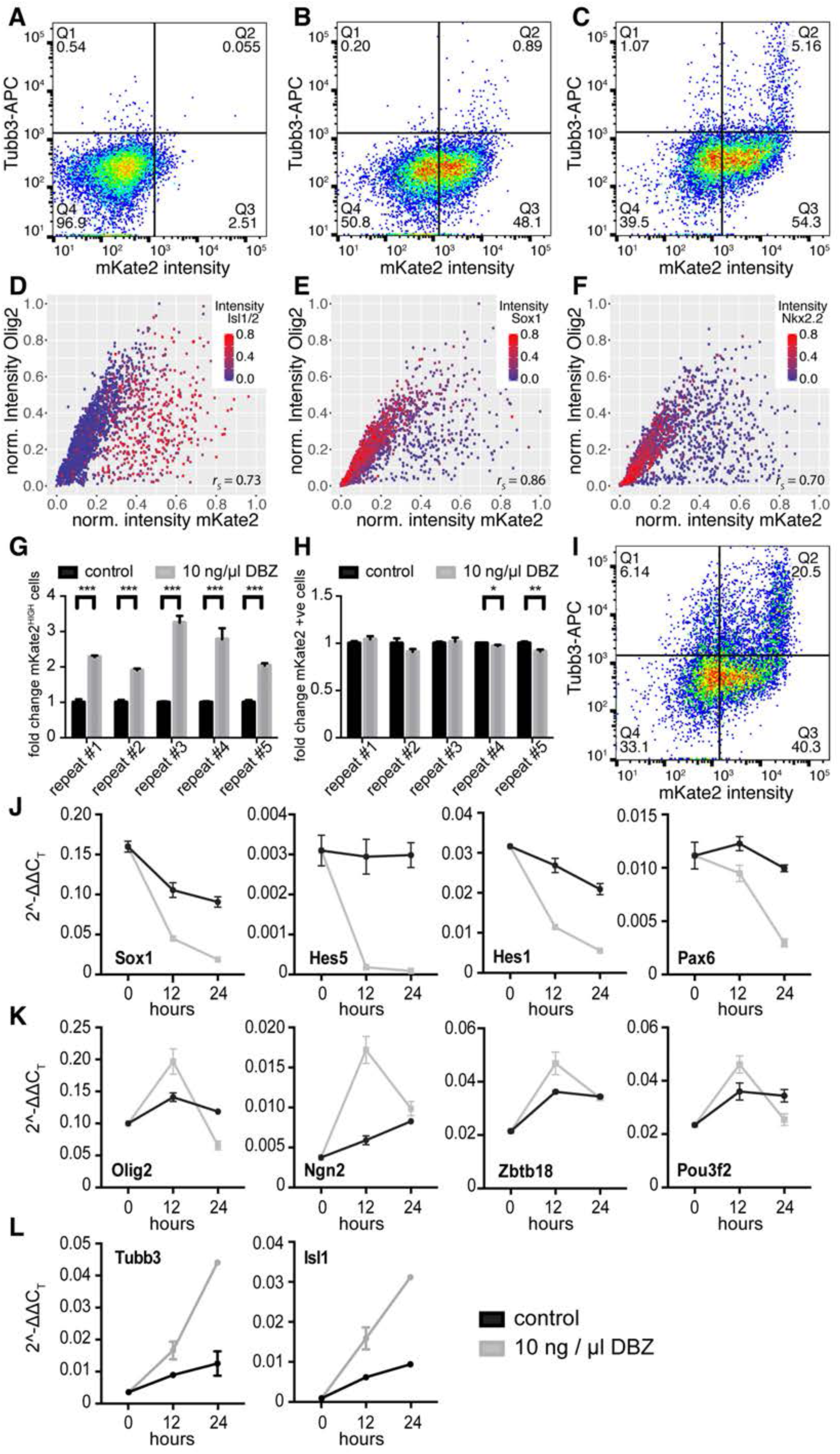
Characterization of the Olig2-mKate2 reporter cell line by flow cytometry and upon Notch inhibition. (A-C) Quantification of mKate2 and Tubb3 fluorescence intensity by flow cytometry at day 4 to day 6 of differentiation. Note that high levels of Tubb3 are predominantly detected in mKate2^HIGH^ cells at day 6 (C). (D-F) Correlation between Olig2 and mKate2 levels in individual nuclei quantified from images in Fig 4C-F. Plots are color coded for levels of Isl1/2 (D), Sox1 (E) and Nkx2.2 (F). (G) Quantification of the fold change in mKate2^HIGH^ cells (see Fig 4L) upon 24 hours Notch inhibition for five experimental repeats by flow cytometry. Each repeat consists of the measurement of three independent dishes for control and Notch inhibition from the same differentiation. *** p < 0.001, unpaired t-test (H) Fold change of mKate2-positive cells (see Fig 4L) upon Notch inhibition (grey) relative to untreated control differentiations (black). Notch inhibition does not cause an overall change in the number of mKate2 positive cells. * p < 0.05; ** p < 0.01, unpaired t-test (I) Quantification of mKate2 and Tubb3 fluorescence intensity by flow cytometry upon 24 hours Notch inhibition. Note that most mKate2^HIGH^ cells differentiated into MNs (compare to Fig S6C). (J-L) RT-qPCR quantification of expression levels of progenitor markers *Sox1*, *Hes5*, *Hes1* and *Pax6* (J), neurogenesis markers *Olig2*, *Ngn2*, *Zbtb18* and *Pou3f2* (K) and MN markers *Tubb3* and *Isl1* (L) after 0, 12 and 24 hours of Notch inhibition (grey) and in untreated controls (black). Note that *Olig2* expression increases in contrast to other progenitor markers after 12 hours of Notch inhibition.

**Fig S7.**
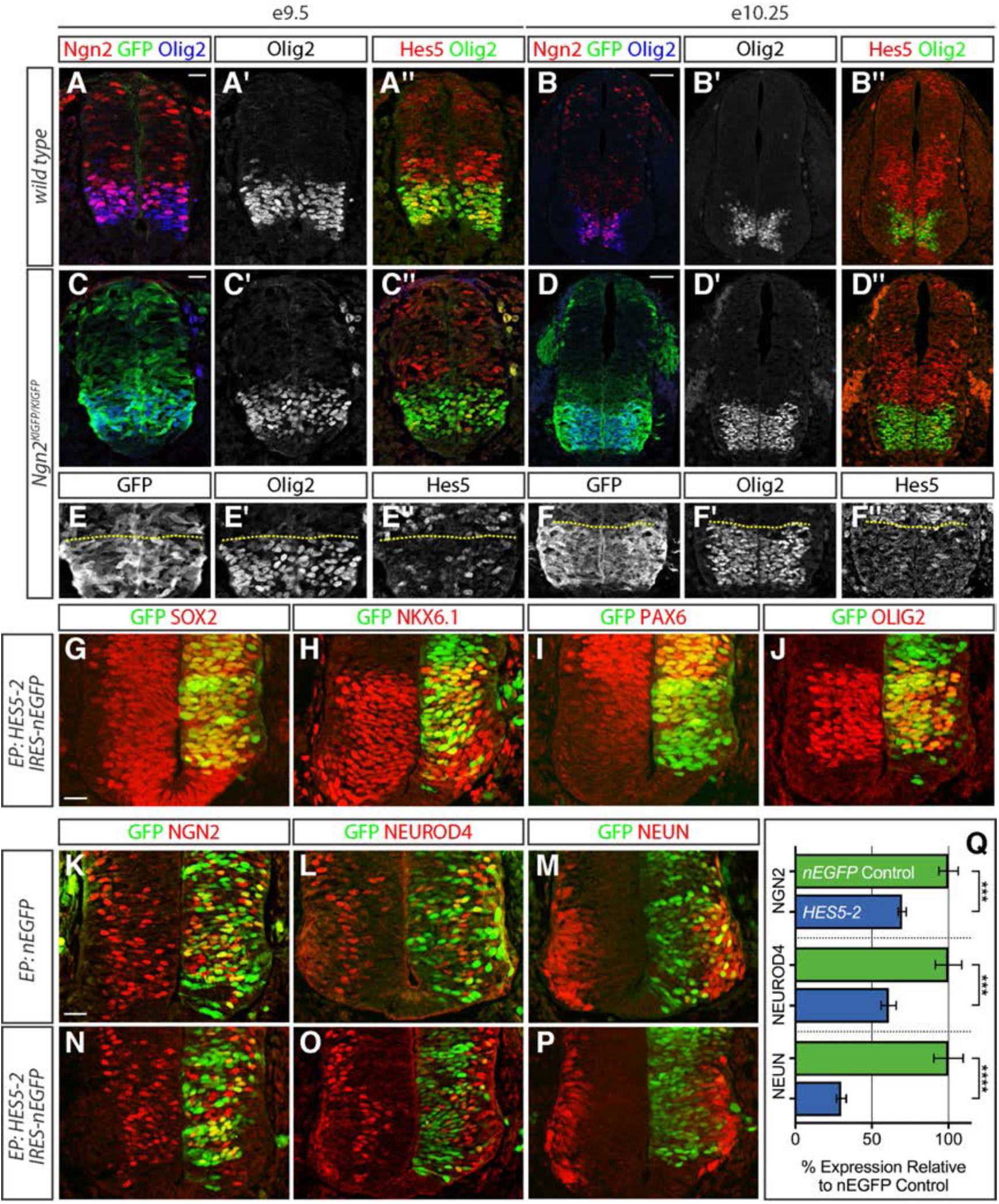
Olig2, not Ngn2, is required for the repression of Hes5, which is necessary for the formation of MNs. (A-D’’) Staining of wild type (A-B’’) and *Ngn2*^*KIGFP*^ mutant (C-D’’) embryonic spinal cords for Olig2, Ngn2, GFP and Hes5 at e9.5 (A,C) and e10.25 (B,D). (E-F’’) GFP expression (E,F) is still increased and Hes5 expression (E’’,F’’) is still reduced in the pMN domain in *Ngn2*^*KIGFP*^ mutant spinal cords (same sections as C-D’’). The yellow dotted line indicates the dorsal boundary of the pMN domain. (G-J) Ectopic expression of cHES5-2 does not affect levels of the progenitor markers SOX2 (G), NKX6.1 (H), PAX6 (I) and OLIG2 (J). (K-P) Ectopic expression of cHES5-2 (N-P) leads to a reduction of NGN2 (K,N), NEUROD4 (L,O) and NEUN (M,P). (K-M) show control electroporations with a nuclear EGFP (nEGFP) expression construct. (Q) Quantification of the effect of ectopic cHES5-2 expression on expression levels of NGN2, NEUROD4 and NEUN relative to nEGFP controls. **** p < 0.0001; *** p < 0.0005, Mann-Whitney test Scale bars = 20 µm (A-A”,C-C”,G-P), 50 µm (B-B”,D-D”).

**Fig S8.**
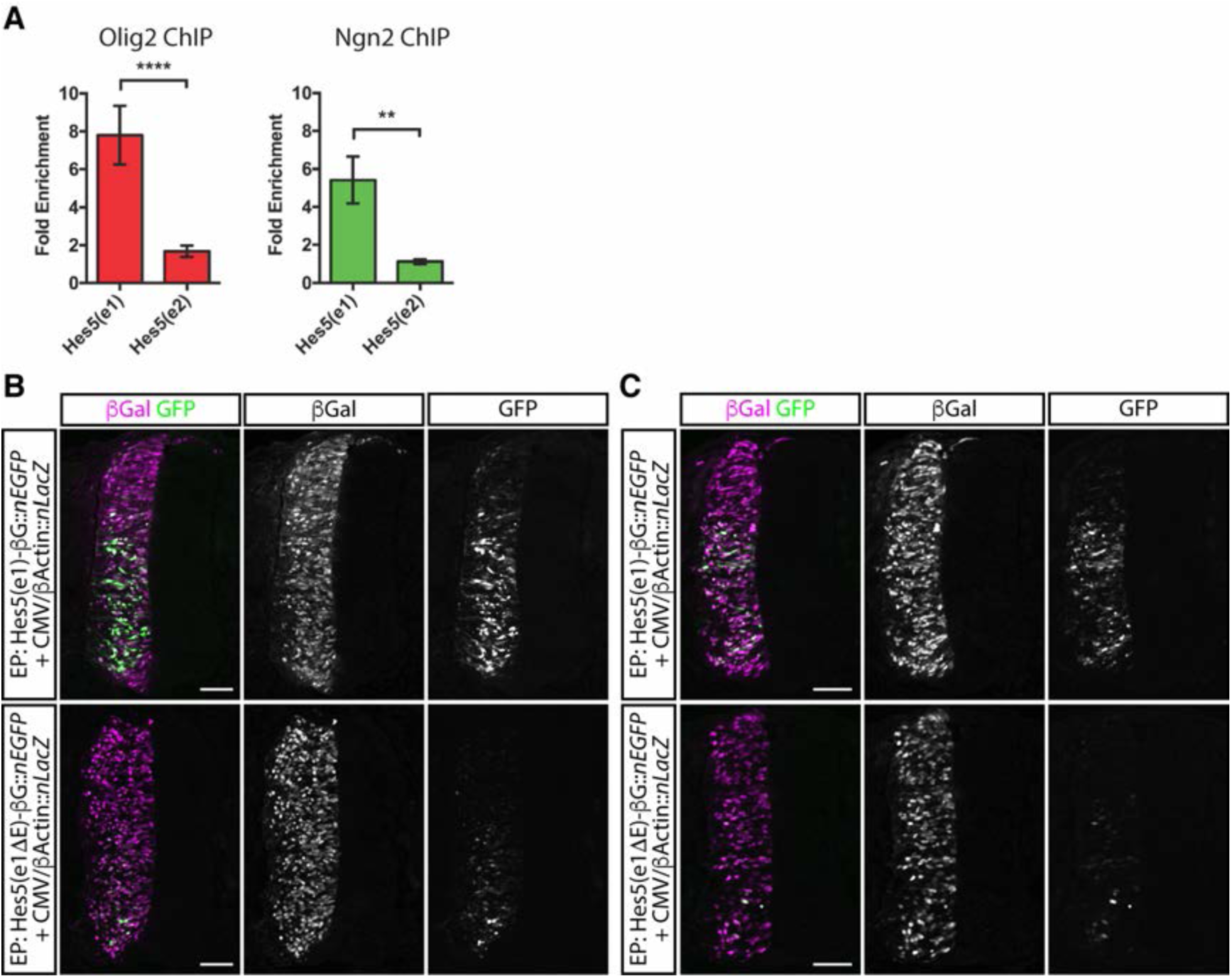
Olig2 and Ngn2 bind to Hes5(e1) in vivo. (A) Both Olig2 and Ngn2 antibodies precipitate the *Hes5(e1)* genomic element from ESC-derived MN progenitors, but not *Hes5(e2),* an unrelated genomic element 3’ to the *Hes5* coding exons that also contains an E-box. Fold enrichment relative to normal rabbit sera or purified IgG is displayed. **** p < 0.0001; ** p < 0.01, Mann-Whitney test. (B,C) Comparison between Hes5(e1)-βG::*nEGFP* (top row) and Hes5(e1βE)-βG::*nEGFP* (bottom row) reporter activities. Sections were imaged using identical settings for each pair. The overall activity of the Hes5(e1βE)-βG::*nEGFP* reporter is lower than that of the Hes5(e1)-βG::*nEGFP* reporter. Scale bars = 50 µm

**Table S1:**
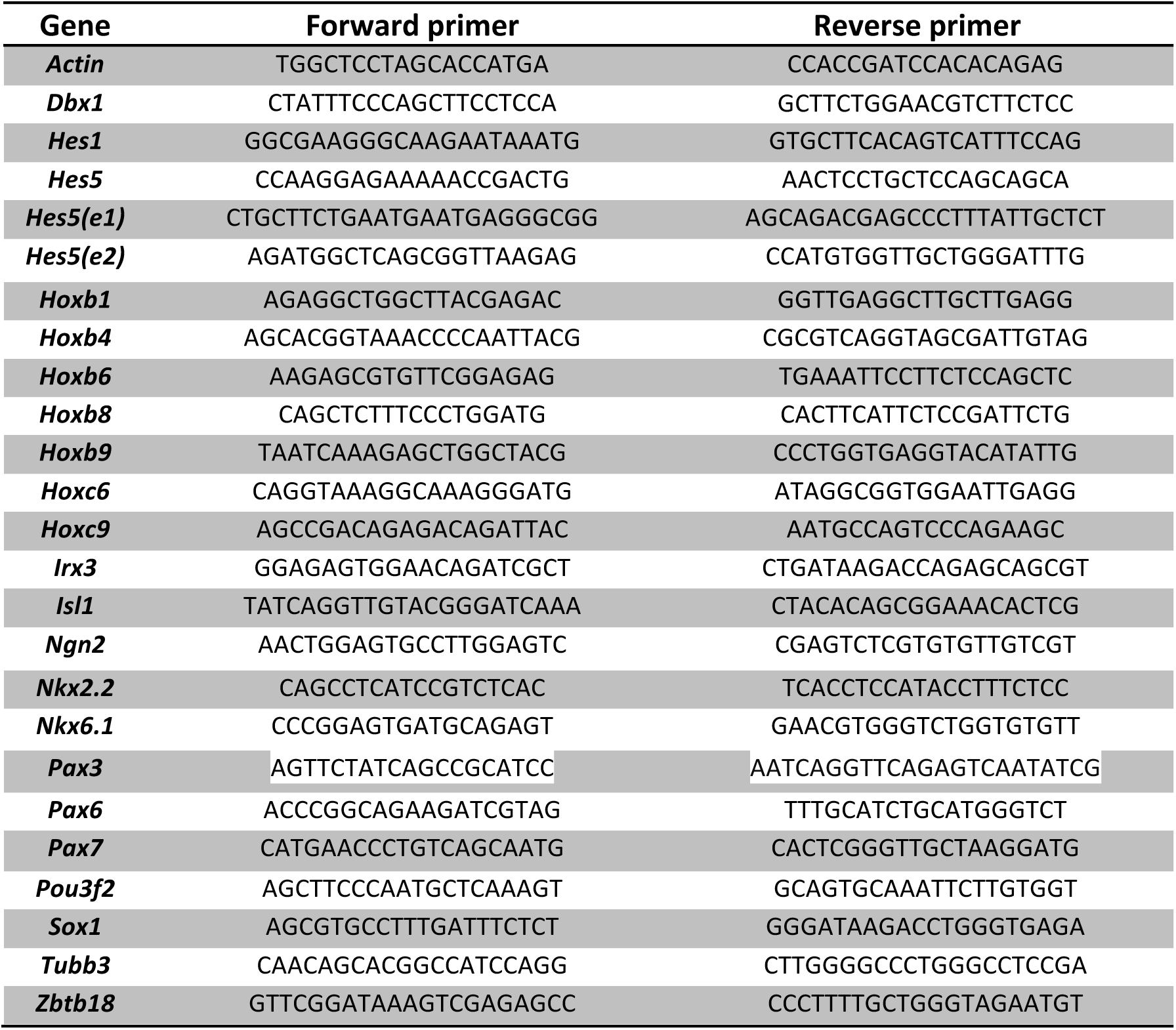
List of qPCR primers.

**Supplementary file 1: Gene modules describing the biological processes represented in the scRNA-seq dataset**

**Supplementary file 2: GO annotations of the 22 gene modules (pval < 0.05)**

## ANALYTICAL SUPPLEMENT

### RNA sequences alignment and pre-processing

Sequences were aligned to the Ensemble mouse genome GRCm38 using Tophat2 (Kim et al., 2013) and counted with HTSeq-count. Cell debris and doublets were removed from the data by inspecting miscroscope images of the microfluidic chips. Low-quality libraries were excluded from the 236 sequenced single-cell transcriptomes if their transcript abundance was less than 10^6^ reads and the number of expressed genes was less than one thousand. The 202 retained libraries (25 cells from day 4, 68 cells from day 5 and 109 cells from day 6) were normalized to read counts per million (CPM). Genes with counts in less than 3 cells or annotated as pseudogenes were excluded from the analysis.

### Cell state identification

To identify the cell states in the dataset, we applied a two-stage strategy aimed at selecting the gene modules demonstrating relevant and concerted patterns of expression. First, we took a data-driven approach to characterize the different modules of interacting genes. From the initial set of 13196 expressed genes, we selected the 2287 genes that showed Spearman correlation (r > 0.4) with at least two other genes. The correlated genes were grouped into 127 gene modules by performing a hierarchical clustering using the Euclidean distance of the z-scored log-transformed gene levels and Ward’s agglomeration criterion (Ward, 1963). The number of modules was selected by determining the “elbow” position in the curve representing the total within-module gene level variation per number of modules. Gene modules were removed according to two criteria: insufficient number of cells expressing the comprised genes and inconsistent gene pattern in these cells. Both criteria were assessed by binarizing gene expression levels using an parameter-free adaptive thresholding method (R function binarize.array from the ArrayBin package). For each cell, we obtained an average expression level per module by averaging the z-scored log-transformed expression levels of all genes belonging to the module. Each of the 127 average expression level distributions were binarized independently. A cell was considered expressing a gene module if the associated Boolean value was true. Modules with fewer than four cells expressing it were excluded. The second criterion was designed to verify that cells expressing a gene module were showing consistently high levels over most of the genes composing the module. We binarized the z-scored log-transformed expression levels of all genes independently. Then, for each module, we calculated the ratio of Boolean values in cells expressing the module (as defined above). We excluded modules where less than half of these Boolean values were true. Twenty-two modules comprising 1064 genes were retained.

Second, functional annotation of the gene modules revealed the global and unbiased description of the biological processes represented in the dataset (see Supplementary files 1 and 2 showing the genes modules and their associated GO terms). In particular, two cell cycle-related gene modules were excluded (Supplementary file 2): gene module 18 containing genes belong to cell cycle phases G2 and M, and primarily associated with the cell division GO term (GO:0051301); and gene module 20, containing G1 and S genes, and associated with the cell cycle GO term (GO:0007049). To focus on cell type characterization, we selected the 10 modules comprising the GO terms associated with embryonic development, i.e. nervous system development (GO:0007399), skeletal system development (GO:0001501), angiogenesis (GO:0001525), cell differentiation (GO:0030154).

### Cell population clustering

In order to define the cell populations present in the dataset, we performed a hierarchical clustering (Ward’s agglomeration criterion) of the Euclidean distances between cells using the z-scored log-transformed expression levels of the 545 genes included in the 10 selected modules (Fig 2A and Fig S3A). The 4 cell clusters containing vascular endothelial and mesodermal cells and the 5 associated gene modules were excluded from the subsequent analysis. 5 gene modules and 306 genes were retained.

### Single-cell state graph

To investigate the dynamical changes of the transcriptional profile as cells differentiate, we developed a method to relate each cell to its closest neighbours in expression space. Unlike cluster analysis which aims to partition cells into groups with similar characteristics, hence breaking the continuity of cell state differentiation, we set out to generate graphs that connect individual cells without requiring the definition of groups. These can reveal the differentiation trajectories and intermediate states that link the clusters of similar cells (the “clustered” populations).

Using the log-transformed expression levels in the 306 genes space, we first calculated the Euclidean distance matrix between each cell and hence constructed a complete weighted graph of cell similarity D. In (Trapnell et al., 2014, Camp et al., 2015), a minimum spanning tree (MST) algorithm was used to extract the subset of cell-cell edges, which forms the backbone of differentiation branches. While MSTs ensure that all cells are connected, they are also sensitive to noise, making the local structure sensitive to small changes in the data (Zemel and Carreira-Perpinan, 2005). To improve robustness to noise of MSTs, we constructed a consensus graph which combines multiple perturbed minimum spanning trees (pMSTs). Each pMST is obtained by calculating a MST from the cell dissimilarity matrix D with a certain ratio j of its elements set to a very large value (j=20%), hence forbidding the recruitment of the associated edges. Individual pMSTs are merged by summing their adjacency matrices into a matrix storing the occurrences of each edge. We then exclude rarely used edges by clustering the non-null edge occurrence distribution using the Fisher method (Fisher,1958) and removing all edges belonging to the first class. This leaves edges that are used repeatedly in multiple permutations and therefore represent good choices for inclusion in MST graphs. The perturb-and-merge algorithm works iteratively until convergence in the number of included edges. The graph visualization shown in Fig 2B,C and Fig S3C were obtained by projecting the graph into 2D where the positions of each cell (node) in the graph were initially random and then adjusted using an iterative force-based layout algorithm, ForceAtlas2 (Jacomy et al., 2014). Gene expression patterns shown in Fig 2C and Fig S3C were smoothed by averaging each cell’s log-transformed gene levels with its neighbors’ log-transformed gene levels. We refer to these transformed levels as “log-smoothed” in the following.

### Pseudo-temporal ordering

One of the advantages of generating a single-cell state graph is the possibility to infer a pseudo-temporal ordering of the gene expression by following the gene expression implied by the spanning tree. The strategy we used was to identify two terminal cell populations, early and late, and then find the K-shortest paths that connect each pair of early and late cells (Martins and Pascoal, 2003). The early population was specified by selecting the 3 cells expressing a combination of highest Irx3 level and lowest Tubb3 level, and the late population by selecting the 3 cells expressing the highest Tubb3 level. A thousand k-shortest paths were generated for each of the 9 pairs of early and late cells. The resulting 9000 paths did not necessarily have the same length, 90.4% of them were formed by between 14 and 17 cells (shortest paths had 13 cells and longest 19 cells). In order to average gene expression along all paths, each of the 9000 paths was rescaled to the same length. Path rescaling was performed by replicating the cell IDs forming a path so that the total rescaled path length would match a constant value set to 41 pseudotime points (Fig 2D). As no path length was a factor of 41, some cell IDs were replicated either 2 or 3 times (13-cell-long paths being the exception with cell IDs replicated 3 or 4 times). For example, 16-cell-long paths had 7 cell IDs repeated 2 times, and 9 cell IDs repeated 3 times. To avoid the introduction of any bias in the repetitions, the choice of replicating a cell ID 2 or 3 times was random. The resulting 9000 equally-sized paths provided a list of 9000 cell IDs for each of the 41 pseudotime points. These lists allowed the calculation of various measurements along the pseudotime scale. In particular, Fig 2E,F and Fig S3E,F show the mean value of the 9000 log-smoothed gene levels for each of the 41 time points. All the pseudo-temporal dynamics were smoothed using a local polynomial regression fit (R function loess with span=0.5).

### Robustness of pseudo-temporal ordering

In order to assess the robustness of our pseudo-temporal orderings, we performed a bootstrapping of the predicted 13 gene expression profiles shown in Fig 2 with 1000 replicates. Following standard bootstrapping procedure (Booth et al., 1993), the cells of each bootstrapped dataset were drawn randomly with replacement. Hence the bootstrapped datasets were composed on average of about 97 different cells while maintaining the original sample size with cells selected multiple times (the expected number of cells selected at least once in a boostrapped dataset is given by 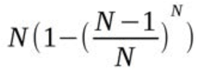 (with N=154 original cells). Following (Haghverdi et al., 2016), we constructed a (1000 by 1000) “self-concordance” matrix for each gene, the elements of which are Spearman correlation of the expression profiles obtained between all pairs of replicates. Calculating the mean and standard deviation of these matrices reads: *Chat* (mean=0.76, sd=0.11), *Hes1* (mean=0.90, sd=0.05), *Hes5* (mean=0.92, sd=0.05), *Irx3* (mean=0.91, sd=0.07), *Isl1* (mean=0.88, sd=0.10), *Isl2* (mean=0.85, sd=0.13), *Lhx3* (mean=0.87, sd=0.12), *Neurod4* (mean=0.88, sd=0.11), *Neurog2* (mean=0.90, sd=0.08), *Nkx6.1* (mean=0.89, sd=0.08), *Olig2* (mean=0.90, sd=0.07), *Pax6* (mean=0.89, sd=0.06), *Tubb3* (mean=0.86, sd=0.09). The percentile confidence intervals for the gene expression profiles are shown in Fig S4.

### Gene variation and dynamical states

Quantification of the metastable states and transition phases were obtained by calculating the global gene variation along pseudotime. To do so, we identified the 2466 genes with higher dispersion, i.e. higher ratio of variance over mean as described in (Satija et al., 2015), and with an average expression level higher than 10 CPM to avoid taking into account low-level gene’s variation. The absolute value of the first derivative of these genes was averaged to define the gene variation (Fig 2D,E,F and Fig S3E,F).

After applying differentiation-and-smoothing twice to gene variation (smoothing with local polynomial regression fit), we obtained a profile showing positive values for periods of higher gene variation and negative values for periods of lower gene variation, hence defining the dynamical states along pseudotime. This operation is equivalent to applying a low-pass Savitzky-Golay filter to the gene variation signal.

